# Adenosylhomocysteinase drives epigenetic and metabolic reprogramming in ZFTA-RELA ependymoma

**DOI:** 10.64898/2026.10.06.757085

**Authors:** Navneesh Yadav, Olivia Pelcher, Raja G.R. Mooli, Robert B. Cameron, Alberto Delaidelli, Kannan B.N. Rao, Bhagyalakshmi Manikandan, Senna Visser, Antonio C.T. Cobos, Vishal Parmar, Yuvraj Kayola, Siva K. Natarajan, Yash Patel, Jiao Zhang, Hardik Shah, Subrahmanya D. Vallabhapurapu, Saifur R. Khan, Aaron Y. Mochizuki, Kalil G. Abdullah, Baoli Hu, Calixto-Hope G. Lucas, Philipp Sievers, Felix Sahm, Andrew M. Donson, Nicholas K. Foreman, Martine F. Roussel, Thomas M. Pearce, Peter Dirks, Kristian W. Pajtler, Stephen C. Mack, Jeremy Rich, Richard J. Gilbertson, Eric C. Holland, Sadeesh K. Ramakrishnan, Taylor J. Abel, Srinivas Ramachandran, Sameer Agnihotri, Bokai Zhu, Sriram Venneti, Brandon Faubert, Michael D. Taylor, Esther Hulleman, Ian F. Pollack, Timothy Phoenix, Kulandaimanuvel Antony Michealraj

## Abstract

Supratentorial ependymomas (ST-EPNs) are aggressive pediatric brain tumors predominantly driven by the recurrent ZFTA-RELA fusion, an oncogenic chromatin regulator that establishes an aberrant transcriptional program. Although metabolic pathways modulate epigenetic regulation by controlling the availability of metabolic cofactors, how ZFTA-RELA coordinates metabolic reprogramming with epigenetic regulation is not well understood. Here, we show that ZFTA-RELA-positive ependymomas exhibit enhanced methionine dependency, elevated adenosyl homocysteinase (AHCY) expression, and epigenetically control aspartate metabolism for tumor growth. Mechanistically, ZFTA-RELA upregulates AHCY, increasing cellular methylation potential which promotes H3K4me3-dependent expression of CAD, the first enzyme complex of de novo pyrimidine biosynthesis, while simultaneously enforcing H3K27me3 mediated repression of ASNS, which encodes asparagine synthetase. Through this coordinated epigenetic program, aspartate utilization is directed toward pyrimidine biosynthesis while endogenous asparagine production is limited. Restricting S-adenosylmethionine availability or genetically or pharmacologically inhibiting AHCY reverses these epigenetic states, leading to CAD downregulation, ASNS upregulation, impaired nucleotide synthesis, and suppression of tumor growth in vitro and in vivo. Similarly, EZH2 inhibition also restores ASNS expression and suppresses tumor proliferation. These findings suggest that ZFTA-RELA EPN maintain ASNS-low phenotype which conditionally make them dependent on extracellular asparagine. We further show that enhanced de novo pyrimidine synthesis not only supports nucleotide production but also provides UDP-sugars required for N-glycosylation and stabilization of the asparagine transporter SLC1A5, thereby promoting extracellular asparagine uptake and tumor proliferation. Collectively, our findings identify AHCY as a critical metabolic regulator of epigenetic and metabolic reprogramming that drives pyrimidine synthesis and establishes extracellular asparagine dependency, revealing actionable metabolic vulnerabilities in ZFTA-RELA-positive ST-EPNs.

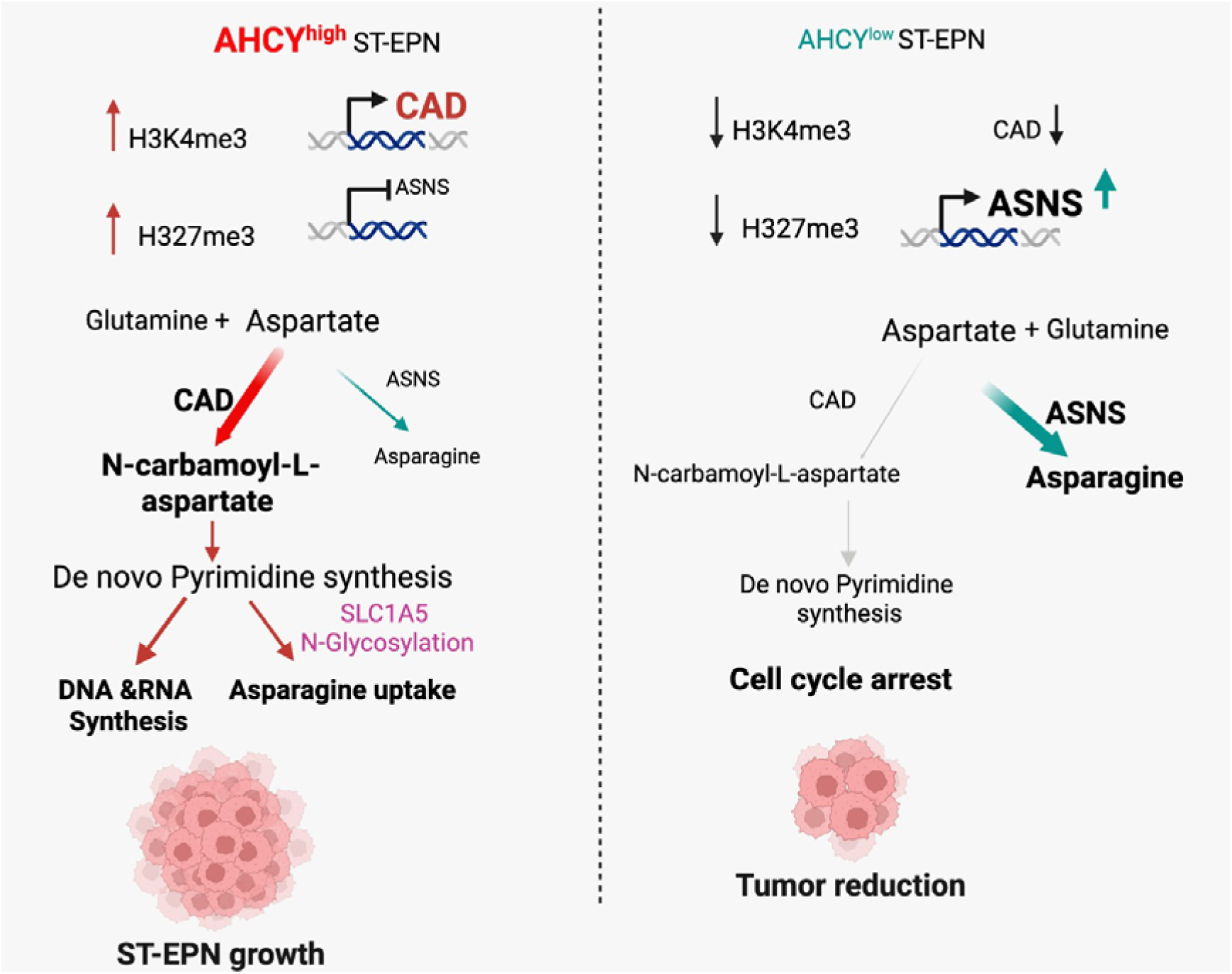

**Key Findings:**

- ZFTA-RELA fusion drives aspartate metabolic reprogramming through chromatin regulation
- Adenosylhomocysteinase (AHCY) epigenetically enhance de novo pyrimidine synthesis and suppress endogenous asparagine synthesis
- ZFTA-RELA EPNs exhibit ASNS**^low^** phenotype and dependence on extracellular asparagine
- Enhanced de novo pyrimidine biosynthesis fuels nucleotide synthesis and SLC1A5-dependent asparagine uptake
- Targeting de novo pyrimidine biosynthesis and depleting extracellular asparagine reveal actionable vulnerabilities in ZFTA-RELA ST-EPN

## Introduction

Ependymomas are malignant tumors that commonly occur in children and comprise molecularly distinct subgroups arising throughout the central nervous system^1,2^. Molecular classification has substantially advanced our understanding of ependymoma biology, revealing genetically defined subgroups, oncogenic mechanisms, epigenetic landscapes, and transcriptional programs^3–7^. Despite these advances, current treatment remains largely limited to maximal safe surgical resection and radiotherapy, with no targeted therapies currently available for ependymoma^8–11^.

Supratentorial ependymomas (ST-EPNs) are primarily driven by recurrent oncogenic fusion events involving either ZFTA or YAP1, with ZFTA-RELA (ZR) fusion-positive tumors representing the predominant subtype (∼70%) and one of the most clinically aggressive forms of ependymoma^2,4,6^. The ZR fusion functions as an aberrant transcriptional regulator that constitutively activates NF-κB signaling and remodels chromatin accessibility to establish tumor-promoting transcriptional programs^12–15^. Recent studies have further demonstrated substantial intratumoral heterogeneity in ST-EPN^16^ and shown that ZR preferentially exploits developmentally accessible chromatin elements, linking its oncogenic activity to the epigenetic state of the potential cell of origin^17^. Additionally, the oncometabolite itaconate has recently been identified as an epigenetic regulator that stabilizes the ZR fusion in ST-EPN^18^.

Metabolic dysregulation is a hallmark of cancer, in which tumor cells reprogram nutrient uptake and metabolism to sustain uncontrolled proliferation and tumor growth^19–21^. Moreover, the availability of metabolic intermediates and cofactors plays a critical role in linking oncogenic mechanism to epigenetic regulation in pediatric brain tumors^22–25^. However, how oncogenic fusion proteins reprogram metabolic pathways and coordinate transcriptional, epigenetic, and downstream oncogenic processes in ZR-positive ST-EPN remains poorly understood.

Here, we demonstrate that ZR-positive ST-EPNs epigenetically reprogram de novo pyrimidine and asparagine biosynthesis through a previously unrecognized mechanism. We identify adenosylhomocysteinase (AHCY), a key methionine-cycle enzyme, as a direct transcriptional target of the ZR fusion. By preventing S-adenosylhomocysteine accumulation, AHCY enhances methylation potential and supports methyltransferase activity, promoting H3K4me3-mediated activation of CAD and H3K27me3-mediated repression of ASNS, thereby channeling aspartate toward de novo pyrimidine biosynthesis. Notably, we identify a unique ASNS-low state in ZR ST-EPN that has not previously been described in pediatric brain tumors, and which makes asparagine a conditionally essential amino acid in these tumors. We further show that de novo pyrimidine biosynthesis not only supports nucleotide production but also generates UDP-sugars required for N-glycosylation-mediated stabilization of SLC1A5, facilitating extracellular asparagine uptake. Consequently, ZR fusion positive tumors become dependent on both de novo pyrimidine biosynthesis and exogenous asparagine. Consistent with these vulnerabilities, L-asparaginase treatment and DHODH inhibition demonstrate significant antitumor activity. Together, our findings identify methionine metabolism drives nucleotide synthesis and asparagine metabolism through non canonical mechanism and reveal these metabolic dependencies as actionable vulnerabilities in ZR-driven ST-EPN.

## Results

### Metabolic regulation in ZFTA-RELA fusion supratentorial ependymoma

To define metabolic dependencies in oncogenic ZFTA-RELA (ZR) fusion driven supratentorial ependymoma (ST-EPN), we leveraged an autochthonous murine ST-EPN model generated by in utero electroporation (IUE) of the developing later lateral ventricles at embryonic day 13.5 with the ZR fusion^26^. Untargeted liquid chromatography-tandem mass spectrometry (LC-MS/MS) metabolomic profiling of murine ZR^Fus^ tumors and adjacent normal cortex/brain tissue (**Fig. 1A**) revealed 132 significantly enriched and 26 depleted metabolites in tumors (**Fig. 1B and Supplementary Table 1**). Pathway analysis of enriched metabolites revealed several significant enriched metabolic programs, including pyrimidine metabolism, methionine metabolism, glycine and serine metabolism, and arginine and proline metabolism (**Fig. 1C**). Independently, we performed LC-MS/MS-based untargeted metabolomic profiling of treatment-naïve surgical specimens from patients with ZR^Fus^ ST-EPN and age-matched normal cortex (**Fig. 1D**). Differential metabolite analysis revealed significantly enriched and depleted metabolites (**Fig. 1E and Supplementary Table 2**). Pathway analysis of enriched metabolites identified methionine metabolism as a significantly enhanced metabolic pathway, followed by glycine and serine metabolism, histidine metabolism, and polyamine biosynthesis (**Fig. 1F**). Integration of human and murine ST-EPN metabolomic datasets revealed consistent enrichment of metabolites associated with methionine metabolism, glycine and serine metabolism, histidine metabolism, and spermidine/spermine biosynthesis in ZR^Fus^ ST-EPN tumors (**Fig. 1G**). Notably, the methionine-cycle metabolite S-adenosylmethionine (SAM) and the methionine salvage pathway intermediate methylthioadenosine (MTA) were among the most significantly elevated metabolites in ZR^Fus^ tumors (**Fig. 1B, E**).

**Fig. 1:**
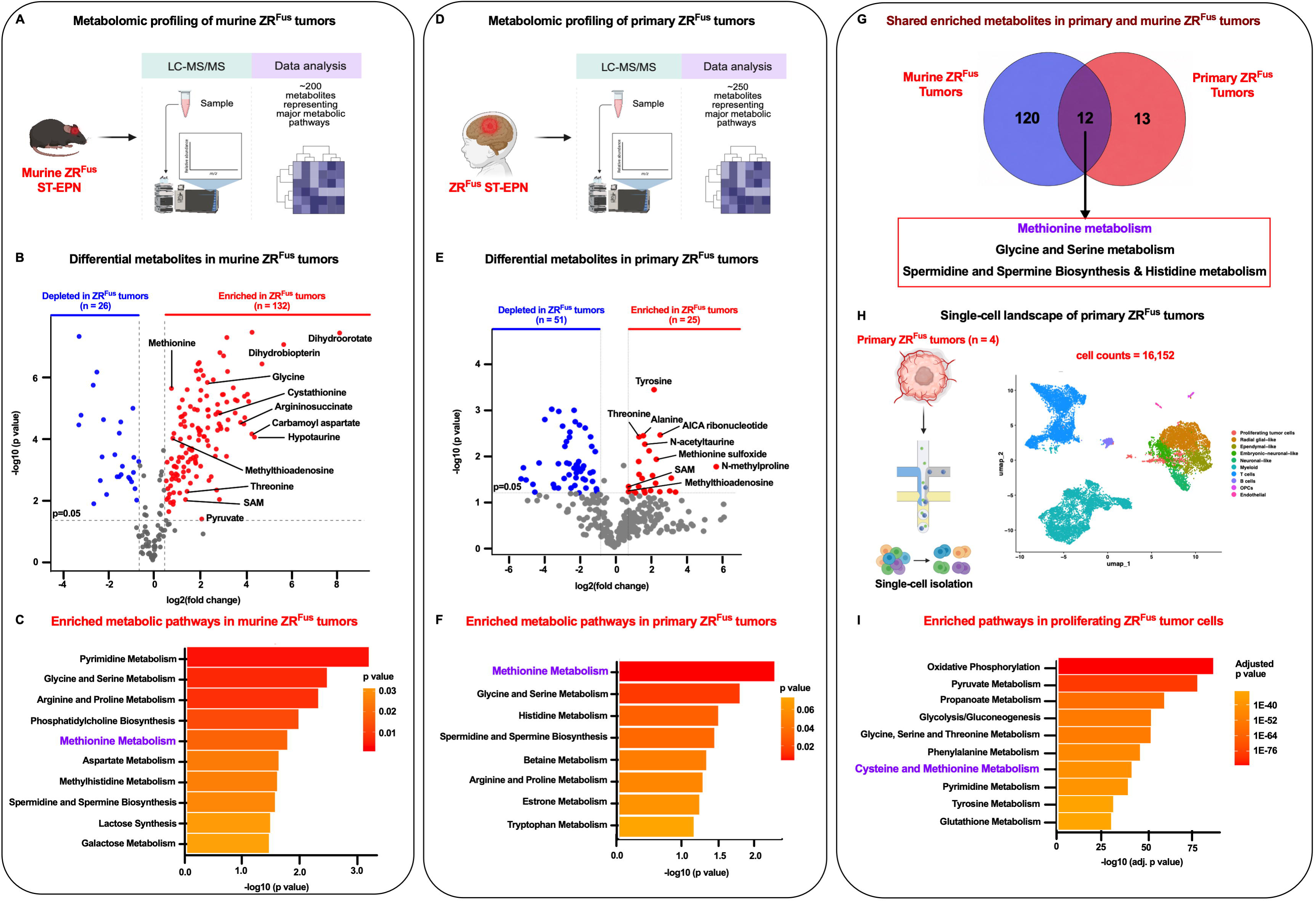
ZFTA-RELA fusion drives metabolic reprogramming in supratentorial ependymoma. (A) Schematic of untargeted liquid chromatography-tandem mass spectrometry (LC-MS/MS) metabolomic profiling of autochthonous murine ZFTA-RELA fusion-positive (ZR^Fus^) supratentorial ependymoma (ST-EPN) tumors. (B) Volcano plot showing differential metabolites in murine ZR^Fus^ tumors compared with adjacent normal cortex. Significantly altered metabolites (p ≤ 0.05 and |log_2_ fold change| ≥ 0.58) are highlighted in red (enriched) and blue (depleted). SAM, S-adenosylmethionine. (C) Top metabolic pathways identified from the 132 metabolites significantly enriched in murine ZR^Fus^ tumors. Methionine metabolism was among the enriched pathways. (D) Schematic of untargeted LC-MS/MS metabolomic profiling of treatment-naïve primary ZR^Fus^ ST-EPN tumors. (E) Volcano plot showing differential metabolites in primary ZR^Fus^ ST-EPN tumors compared with normal cortex. Significantly altered metabolites (p ≤ 0.05 and |log_2_ fold change| ≥ 0.58) are highlighted in red (enriched) and blue (depleted). (F) Top metabolic pathways identified from the 25 metabolites significantly enriched in primary ZR^Fus^ tumors. Methionine metabolism was the most significantly enriched pathway. (G) Venn diagram showing shared enriched metabolites between primary human and murine ZR^Fus^ tumors. The 12 commonly enriched metabolites belonged to methionine metabolism, glycine and serine metabolism, spermidine and spermine biosynthesis, and histidine metabolism. (H) Schematic of the single-cell RNA sequencing (scRNA-seq) analysis (left) and Uniform Manifold Approximation and Projection (UMAP) visualization of 16,152 cells from primary ZR^Fus^ ST-EPN tumors (right). Each dot represents an individual cell and is color-coded according to cell identity. OPCs, oligodendrocyte progenitor cells. (I) Top enriched pathways in proliferating ZR^Fus^ tumor cells identified by scRNA-seq differentially expressed genes. Pathways with adjusted p ≤ 0.05 were considered significant. Cysteine and methionine metabolism was among the enriched metabolic pathways. For panels B and E, differential metabolite analysis was performed using four and three biological replicates per group, respectively. Panel H comprises scRNA-seq data from four primary ZR^Fus^ tumors.

To determine the metabolic pathways that are regulated by oncogenic ZR fusion, we established isogenic patient derived ST-EPN models with and without ZR fusion expression using CRISPR/Cas9-mediated ZR fusion-specific guide RNA (**Extended Data Figs. 1A-D**). Relative to ZR fusion-positive cells, matched ZR fusion knockout cells exhibited significant reductions in metabolites associated with methionine metabolism, pyrimidine metabolism, glutathione metabolism, and pyruvate metabolism (**Extended Data Fig. 1E and Supplementary Table 3**). Together, metabolic profiling of ZR^Fus^ EPN in vivo and in vitro models demonstrated broad metabolic reprogramming with amino acid metabolism, particularly pinpointing methionine metabolism as an important metabolic dependency.

Given that ZR fusion functions as an aberrant transcription factor and chromatin regulator^6,7,27^, and can regulate transcriptional programs associated with metabolic pathways, we next examined metabolic gene expression and pathway enrichment. To this end, we analyzed single-cell RNA sequencing (scRNA-seq) dataset from patient derived ZR^Fus^ tumors (**Fig. 1H**). KEGG pathway enrichment analysis^28^ revealed that proliferating tumor cells exhibited increased expression of genes associated with methionine and cysteine metabolism and pyrimidine metabolism (**Fig. 1I, Extended Data Fig. 1F, and Supplementary Tables 4, 5**). Similarly, transcriptomic analyses of ZR fusion isogenic patient-derived ST-EPNs models (**Extended Data Figs. 2A-C and Supplementary Tables 6, 7**) and murine neural stem cells (mNSCs) ectopically expressing the ZR fusion (**Extended Data Figs. 2D-H and Supplementary Table 8**) revealed that genes involved in the methionine metabolism, one-carbon metabolism, pyrimidine, purine metabolism, and glycolysis/gluconeogenesis were upregulated in a ZR fusion-dependent manner (**Extended Data Figs. 1I-K**).

Collectively, our integrated metabolomic and transcriptomic analyses across human tumors, genetically engineered mouse models, and isogenic models identified key metabolic pathways regulated by ZR fusion and highlight methionine metabolism as one of the critical metabolic dependencies in ZR fusion driven ST-EPN. Given that methionine is an essential amino acid and critical for downstream metabolic and various cellular functions, we investigated the mechanism underlying this methionine dependency.

### ZFTA-RELA fusion demonstrates enhanced methionine dependency

Metabolic investigations using ZR^Fus^ ST-EPN models consistently identified methionine metabolism as a critical metabolic dependency. Tumor cells require methionine to support rapid and sustained proliferation^19,29,30^. Besides being essential for protein synthesis, methionine is also metabolized into downstream intermediates that fuel cellular metabolism and methylation-dependent processes (**Fig. 2A**). Most standard cell culture media contain supraphysiological concentrations of methionine, typically ranging between 200 and 300 μM. In contrast, physiological serum methionine concentrations have been estimated to range from approximately 3 to 30 μM^31^. Consistent with its essential role, methionine can support protein synthesis even at low extracellular concentrations, with 1.39 μM reported to sustain half-maximal acute protein synthesis^32^. We therefore sought to determine how physiologically relevant methionine availability influenced ZR^Fus^ tumor growth. Using limiting dilution assays (LDAs), we examined sphere-forming capacity under physiologically relevant methionine concentrations. We found that 30 μM methionine significantly enhanced sphere-forming potential in patient-derived ZR^Fus^ ST-EPN models, as well as in mNSCs expressing ZR fusion, whereas sphere formation was markedly reduced under 5 μM methionine conditions (**Figs. 2B, C**). Conversely, methionine availability of 5 μM or 30 μM had no significant effect on sphere formation in neural stem cells transduced with an empty vector (**Fig. 2C**), suggesting that ZR^Fus^ tumors exhibit heightened dependence on physiological methionine levels to sustain tumor cell growth and self-renewal. Methionine dependency was further supported by intracellular methionine levels being significantly higher in ZR^Fus^ tumors compared with adjacent normal cortex (**Fig. 2D**) and with the ZFTA-MAML3 ST-EPN subtype (**Fig. 2E**). Enhanced methionine uptake exhibited correlation with expression of high-affinity methionine transporters SLC7A5^33,34^ and SLC43A2^35,36^ in murine ZR^Fus^ ST-EPN tumors (**Figs. 2F, G**) and in vitro models (**Fig. 2H**). Consistent with this, scRNA-seq of IUE-derived murine ZR fusion ST-EPN tumors also demonstrated enriched expression of *Slc7a5* and *Slc43a2* within malignant cell populations (**Extended Data Fig. 3A**).

**Fig. 2:**
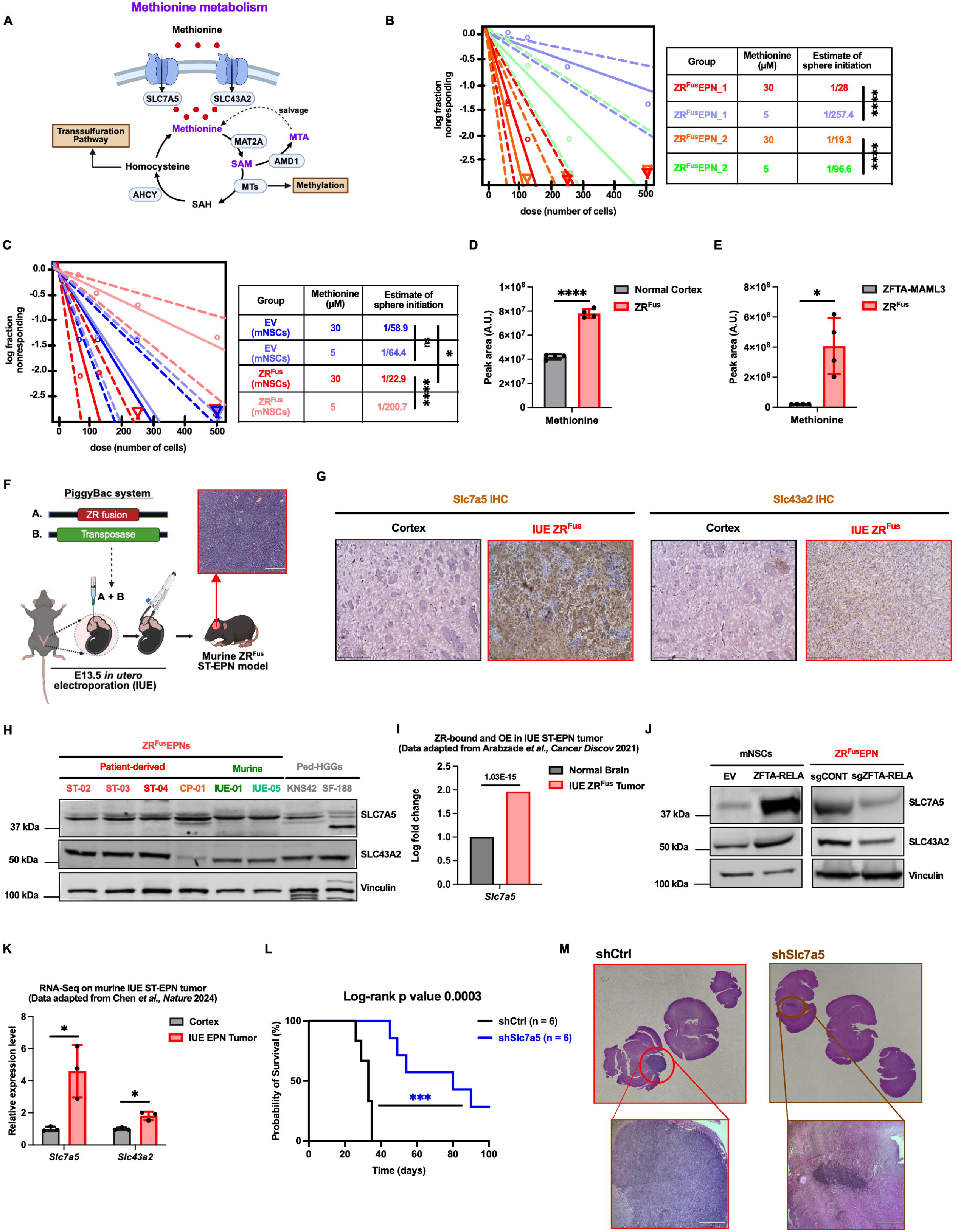
ZFTA-RELA fusion promotes methionine dependence through upregulation of methionine transporters. (A) Schematic of methionine metabolism illustrating methionine uptake through the transporters SLC7A5 and SLC43A2, the methionine cycle, S-adenosylmethionine (SAM)-dependent methylation, the methionine salvage pathway, and the transsulfuration pathway. SAH, S-adenosylhomocysteine; MTA, methylthioadenosine; MAT2A, methionine adenosyltransferase 2A; AHCY, adenosylhomocysteinase; AMD1, adenosylmethionine decarboxylase 1; MTs, methyltransferases. (B) Extreme limiting dilution assay (ELDA) of two patient-derived ZFTA-RELA fusion-positive (ZR^Fus^) supratentorial ependymoma (ST-EPN) cell lines cultured under physiological (30 μM) or methionine-restricted (5 μM) conditions. (C) ELDA of mouse neural stem cells (mNSCs) expressing empty vector (EV) or ZFTA-RELA cultured under physiological (30 μM) or methionine-restricted (5 μM) conditions. (D) Relative abundance of methionine in murine ZR^Fus^ tumors compared with normal cortex. (E) Relative abundance of methionine in patient-derived ZR^Fus^ cells compared with ZFTA-MAML3 fusion-positive ST-EPN cells. (F) Schematic of the PiggyBac-based in utero electroporation (IUE) model used to generate murine ZR^Fus^ ST-EPN tumors at embryonic day 13.5 (E13.5). Scale bar, 220 μm. (G) Representative immunohistochemistry (IHC) images showing SLC7A5 and SLC43A2 expression in normal murine cortex and IUE-derived ZR^Fus^ tumors. Scale bar, 200 μm. (H) Representative immunoblots showing expression of the methionine transporters SLC7A5 and SLC43A2 in patient-derived and IUE-derived murine ZR^Fus^ cell lines compared with pediatric high-grade glioma (Ped-HGG) cell lines. Vinculin served as the loading control. (I) Relative expression of *Slc7a5* in IUE-derived murine ZR^Fus^ tumors compared with normal brain. Data were obtained from the ZFTA-RELA-bound and transcriptionally upregulated gene dataset reported by Arabzade *et al.* (*Cancer Discovery*, 2021). (J) Representative immunoblots showing expression of SLC7A5 and SLC43A2 in mNSCs expressing EV or ZFTA-RELA and in patient-derived ZR^Fus^ cells expressing control single-guide RNA (sgRNA; sgCONT) or sgRNA targeting ZFTA-RELA (sgZFTA-RELA). Vinculin served as the loading control. (K) Relative expression of *Slc7a5* and *Slc43a2* in IUE-derived murine ZR^Fus^ tumors compared with normal brain. Data were obtained from the RNA sequencing (RNA-seq) dataset reported by Chen *et al.* (*Nature*, 2024). (L) Kaplan-Meier survival analysis of immunocompetent C57BL/6J mice bearing orthotopic IUE-derived murine ZR^Fus^ tumors expressing control shRNA (shCtrl) or shRNA targeting Slc7a5 (shSlc7a5). (M) Representative hematoxylin and eosin (H&E)-stained coronal brain sections from mice bearing orthotopic shCtrl or shSlc7a5 tumors. Insets show higher-magnification views of the tumor regions. Scale bar, 220 μm.For panels D and E, data are presented as mean ± SD from four independent samples per group. For panel K, data are presented as mean ± SD from three independent samples per group. For panels B and C, stem cell frequencies were estimated using ELDA from eight replicate wells per cell dose, and differences between groups were assessed using the ELDA likelihood ratio test. Panel L includes six mice per group. Statistical significance was determined using an unpaired two-tailed t-test with Welch’s correction for panels D, E, and K and the log-rank test for panel L. *p < 0.05, ***p < 0.001, ****p < 0.0001; ns, not significant.

Next, we investigated whether the ZR fusion directly regulated methionine transporter expression. Analysis of previously published genome-wide ZR binding datasets identified *Slc7a5* as a direct transcriptional target that is upregulated in ST-EPN tumors^27^ (GEO: GSE161679) (**Fig. 2I and Extended Data Fig. 3B**) and immunoblot analysis of isogenic models suggest that SLC7A5 expression was regulated in a ZR fusion dependent manner, whereas SLC43A2 showed less robust changes in protein expression (**Fig. 2J**). This observation was further supported by analysis of a recently published RNA-seq dataset from an IUE-derived ZR ST-EPN model^37^ (GEO: GSE246033), in which *Slc7a5* expression was markedly increased in tumors relative to normal cortex, while *Slc43a2* showed a more modest increase (**Fig. 2K**). Based on the increased methionine dependency of ST-EPNs and the fusion-dependent regulation of SLC7A5, we functionally assessed its role in tumor maintenance of ST-EPN. Knockdown of Slc7a5 in murine ST-EPN allograft model significantly prolonged survival of tumor-bearing mice (**Figs. 2L, M**), establishing SLC7A5 as a critical metabolic candidate of methionine uptake and tumor growth in ZR fusion driven ST-EPN.

### ZFTA-RELA reprograms methionine metabolism through adenosylhomocysteinase (AHCY)

To determine how the methionine cycle is regulated in ST-EPN, methionine-cycle intermediates were quantified. Especially SAM, the principal methyl donor for cellular methylation reactions, was significantly elevated in ZR^Fus^ ST-EPN compared with normal brain tissue and ZFTA-MAML3 ST-EPN sub-type (**Fig. 3A and Extended Data Fig. 3C**). In contrast, S-adenosylhomocysteine (SAH) levels remained unchanged, resulting in a marked increase in the SAM/SAH ratio in murine tumors (**Fig. 3A**). Consistent with these findings, patient-derived ZR^Fus^ ST-EPN tumors also exhibited significantly higher SAM/SAH ratio than normal human brain tissue and posterior fossa A (PFA) ependymomas (**Fig. 3B**).

**Fig. 3:**
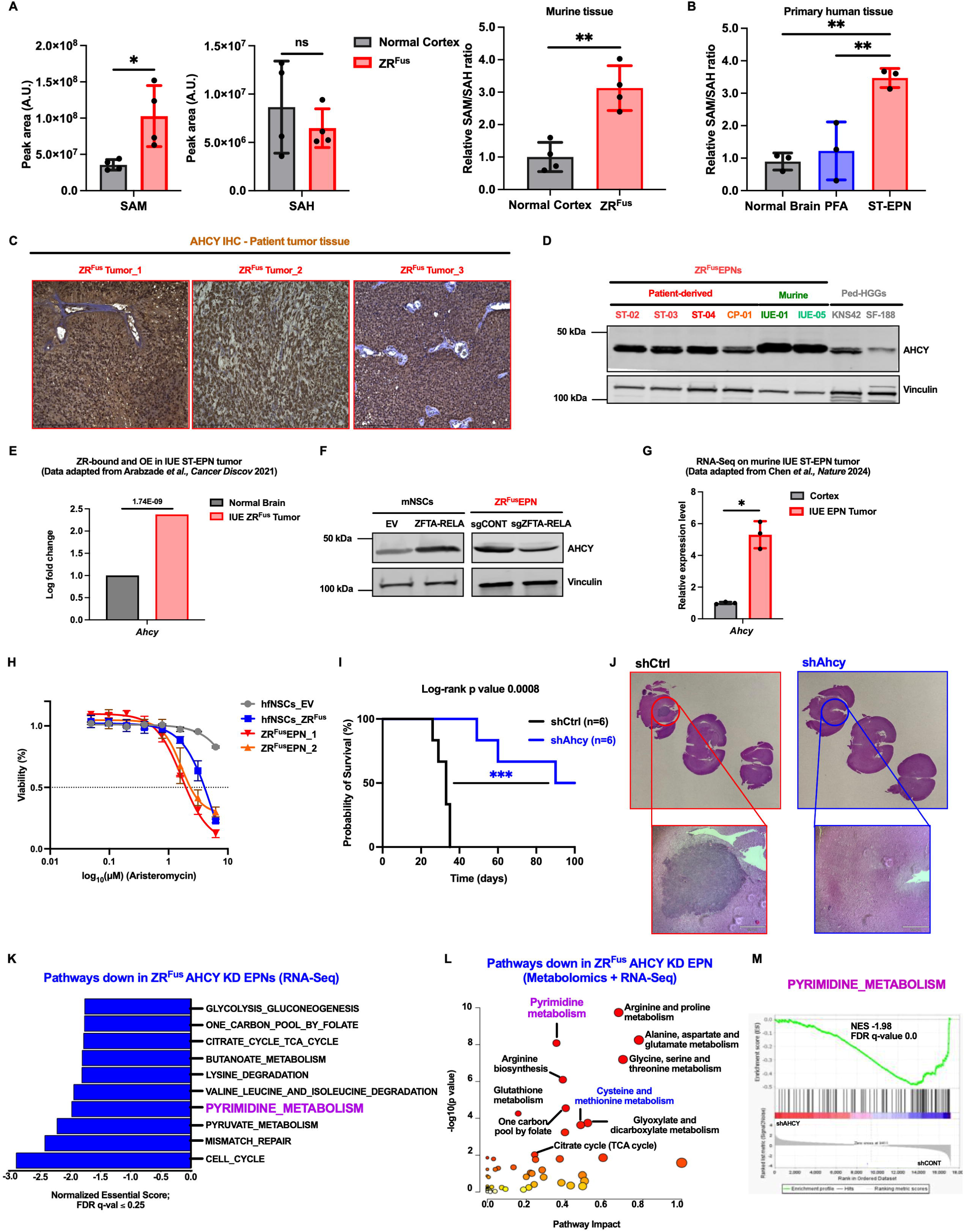
AHCY supports ZFTA-RELA fusion tumor growth and pyrimidine metabolism. (A) Relative abundance of S-adenosylmethionine (SAM) and S-adenosylhomocysteine (SAH), and the relative SAM/SAH ratio, in murine ZFTA-RELA fusion-positive (ZR^Fus^) tumors compared with normal cortex. (B) Relative SAM/SAH ratio in normal brain, posterior fossa A (PFA) ependymomas, and primary ZR^Fus^ supratentorial ependymoma (ST-EPN) tumors. (C) Representative immunohistochemistry (IHC) images showing AHCY expression in three primary ZR^Fus^ ST-EPN tumors. Scale bar, 200 μm. (D) Representative immunoblots showing AHCY expression in patient-derived and in utero electroporation (IUE)-derived murine ZR^Fus^ cell lines compared with pediatric high-grade glioma (Ped-HGG) cell lines. Vinculin served as the loading control. (E) Relative expression of *Ahcy* in IUE-derived murine ZR^Fus^ tumors compared with normal brain. Data were obtained from the ZFTA-RELA-bound and transcriptionally upregulated gene dataset reported by Arabzade *et al.* (*Cancer Discovery*, 2021). (F) Representative immunoblots showing AHCY expression in mouse neural stem cells (mNSCs) expressing empty vector (EV) or ZFTA-RELA and in patient-derived ZR^Fus^ cells expressing control single-guide RNA (sgRNA; sgCONT) or sgRNA targeting ZFTA-RELA (sgZFTA-RELA). Vinculin served as the loading control. (G) Relative expression of *Ahcy* in IUE-derived murine ZR^Fus^ tumors compared with normal cortex. Data were obtained from the RNA sequencing (RNA-seq) dataset reported by Chen *et al.* (*Nature*, 2024). (H) Dose-response curves showing viability of human fetal neural stem cells (hfNSCs) expressing EV or ZFTA-RELA and two patient-derived ZR^Fus^ cell lines following 72 h treatment with increasing concentrations of the AHCY inhibitor aristeromycin. (I) Kaplan-Meier survival analysis of immunocompetent C57BL/6J mice bearing orthotopic IUE-derived murine ZR^Fus^ tumors expressing control shRNA (shCtrl) or shRNA targeting Ahcy (shAhcy). (J) Representative hematoxylin and eosin (H&E)-stained coronal brain sections from mice bearing orthotopic shCtrl or shAhcy tumors. Insets show higher-magnification views of the tumor regions. Scale bar, 220 μm. (K) Top downregulated pathways identified by Gene Set Enrichment Analysis (GSEA) following AHCY knockdown (KD). RNA-seq data from two independent patient-derived ZR^Fus^ cell lines were combined for pathway analysis. Pathways with false discovery rate (FDR) q ≤ 0.25 were considered significant. (L) Integrated metabolomic and transcriptomic pathway analysis following AHCY KD in patient-derived ZR^Fus^ cells, incorporating significantly depleted metabolites (p ≤ 0.05 and log₂ fold change ≤ −1) and downregulated genes (adjusted p ≤ 0.05 and log₂ fold change ≤ −1). Pyrimidine metabolism was among the suppressed pathways. (M) GSEA enrichment plot showing significant suppression of pyrimidine metabolism following AHCY KD (normalized enrichment score (NES) = −1.98, FDR q = 0.0). For panels A and B, data are presented as mean ± SD from four and three biological replicates per group, respectively. For panel G, data are presented as mean ± SD from three independent samples per group. For panel H, data are presented as mean ± SD from three independent samples per condition. For panels K and M, RNA-seq was performed using three independent samples per group. For panel L, metabolomic and transcriptomic analyses were performed using three independent samples per group. Panel I includes six mice per group. Statistical significance was determined using an unpaired two-tailed t-test with Welch’s correction for panels A and G, one-way ANOVA with Šidák’s multiple-comparisons test for panel B, and the log-rank test for panel I. *p < 0.05, **p < 0.01, ***p < 0.001; ns, not significant.

SAM is generated from methionine through the activity of methionine adenosyltransferase 2A (MAT2A). After donating its methyl group in methylation reactions, SAM is converted to SAH, which is then hydrolyzed to homocysteine and adenosine^29^ (**Fig. 2A**). SAH is also a potent competitive inhibitor of methyltransferases that occupies the same catalytic binding pocket as SAM and suppresses methyltransferase activity when accumulated^38,39^. Thus, while increased SAM synthesis enhances methyl-donor availability, sustained methylation potential additionally requires efficient clearance of inhibitory SAH. The observation that ZR^Fus^ ST-EPN tumors exhibited elevated methionine and SAM levels without a corresponding increase in SAH suggested the presence of an active mechanism preventing SAH accumulation. This led to investigation of adenosylhomocysteinase (AHCY), the enzyme responsible for hydrolyzing SAH to homocysteine^40^, and previously implicated as a critical metabolic regulator in multiple cancers^41,42^. We found that AHCY expression was consistently elevated in patient ZR^Fus^ ST-EPN tumors (**Fig. 3C**), as well as in corresponding proliferative cell line models (**Fig. 3D**). Supporting this observation, analysis of previously published dataset^27^ (GEO: GSE161679) confirmed elevated *Ahcy* expression in murine ST-EPN tumors and identified *Ahcy* as a direct transcriptional target of the ZR fusion (**Fig. 3E**). This was further supported by ZR fusion isogenic models, which demonstrated that AHCY upregulation is ZR fusion-dependent (**Fig. 3F**), and by a published RNA-seq dataset from an IUE-derived ZR ST-EPN model^37^ (GEO: GSE246033), in which *Ahcy* expression was markedly increased in tumors relative to normal cortex (**Fig. 3G**). Functional studies revealed that genetic suppression of AHCY using targeted shRNA, or pharmacological inhibition of its enzymatic activity using aristeromycin^43^, profoundly impaired proliferation of ZR^Fus^ ST-EPN cells (**Fig. 3H and Extended Data Fig. 3D**). Likewise, isogenic neural stem cell models expressing the ZR fusion displayed increased sensitivity to AHCY inhibition, indicating a selective dependence on AHCY activity (**Fig. 3H**). Importantly, AHCY knockdown in a murine ZR-fusion ependymoma model significantly reduced tumor burden and prolonged survival (**Figs. 2I, J**), demonstrating a functional role for AHCY in ZR fusion-driven tumor growth.

Because AHCY generates homocysteine, which can subsequently enter the transsulfuration pathway or be recycled back to methionine^29^, we investigated whether restoration of this downstream metabolite could rescue the effects of AHCY inhibition. Supplementation of AHCY inhibitor-treated ZR^Fus^ ST-EPN cells with exogenous homocysteine failed to restore cellular proliferation (**Extended Data Fig. 3E**). This suggests that the essential function of AHCY in ST-EPN is not merely the production of homocysteine and driving methionine cycle forward but also required for effective clearance of SAH and maintains more SAM to SAH ratio which facilitate robust methyltransferase activity. Therefore, we investigated AHCY regulated transcriptional and metabolic programs in ZR^Fus^ ST-EPNs. Differential gene-expression analysis revealed significant perturbations in pathways involved in cell-cycle progression, pyrimidine metabolism, pyruvate metabolism, tricarboxylic acid (TCA) cycle function, one-carbon metabolism, and glycolysis/gluconeogenesis (**Fig. 3K and Extended Data Figs. 3F, G**). Untargeted metabolomic profiling of control and AHCY knockdown cells, followed by integrated analysis with transcriptomic datasets, identified significant depletion of genes and metabolites associated with pyrimidine metabolism, alanine/aspartate/glutamate metabolism, arginine and proline metabolism, and cysteine/methionine metabolism (**Figs. 3L, M and Extended Data Fig. 3H and Supplementary Tables 9, 10**). Consistent with these observations, integrated metabolomic and transcriptomic profiling of patient-derived fusion knockout models and murine ZR fusion tumors identified de novo pyrimidine biosynthesis as a prominent metabolic program activated downstream of the ZR fusion (**Extended Data Fig. 3I, J**). These results suggest that ZR fusion-driven AHCY expression promotes a broader metabolic program in ST-EPN that extends beyond methionine metabolism to support anabolic pathways required for tumor growth. Notably, a recent multi-omics study reported that high AHCY expression was associated with poor clinical outcome and increased expression of pyrimidine metabolic genes in bladder cancer^42^, although the mechanistic basis for this association remains unclear. This prompted us to investigate the molecular mechanisms by which AHCY promotes de novo pyrimidine synthesis in ZR fusion-driven ST-EPN.

### AHCY drives de novo pyrimidine biosynthesis through aspartate metabolic regulation

Aspartate serves as an essential substrate for de novo pyrimidine synthesis and asparagine biosynthesis. Together with glutamine-derived amide nitrogen, aspartate is utilized by CAD to generate N-carbamoyl-L-aspartate, the first committed intermediate of de novo pyrimidine biosynthesis, while endogenous asparagine-producing enzyme ASNS also use glutamine-derived amide nitrogen and catalyzes the conversion of aspartate to asparagine (**Fig. 4A**). We found that loss of AHCY in ZR^Fus^ EPN cells resulted in significant reductions in aspartate and the key de novo pyrimidine biosynthesis intermediates, N-carbamoyl-L-aspartate and dihydroorotate, whereas asparagine accumulated concomitantly (**Figs. 4B, C and Supplementary Table 9**). Significant accumulation of asparagine but relative reduction in aspartate and N-carbamoyl-L-aspartate suggests previously unrecognized metabolic regulation by AHCY. While aspartate-generating enzyme glutamic-oxaloacetic transaminase 2 (GOT2) expression was unaffected by AHCY depletion, expression of aspartate dependent biosynthetic enzyme CAD was significantly reduced (**Fig. 4D**). Conversely, ASNS, which competes with CAD for aspartate was significantly increased following AHCY depletion (**Figs. 4D, E**). The correlation of enzymes expression and metabolomic alterations observed in AHCY-depleted cells suggest that AHCY fine-tunes aspartate utilization for nucleotide biosynthesis while restricting its availability for endogenous asparagine production. Importantly, we found downregulation of AHCY did not alter CAD and ASNS expression in ZFTA-MAML3 ST-EPN cells (**Fig. 4F and Supplementary Tables 10,11**) and pediatric high-grade glioma models (**Extended Data Fig. 3K**), suggesting that ZR^Fus^ tumors selectively exhibit AHCY-dependency for regulation of aspartate metabolism.

**Fig. 4:**
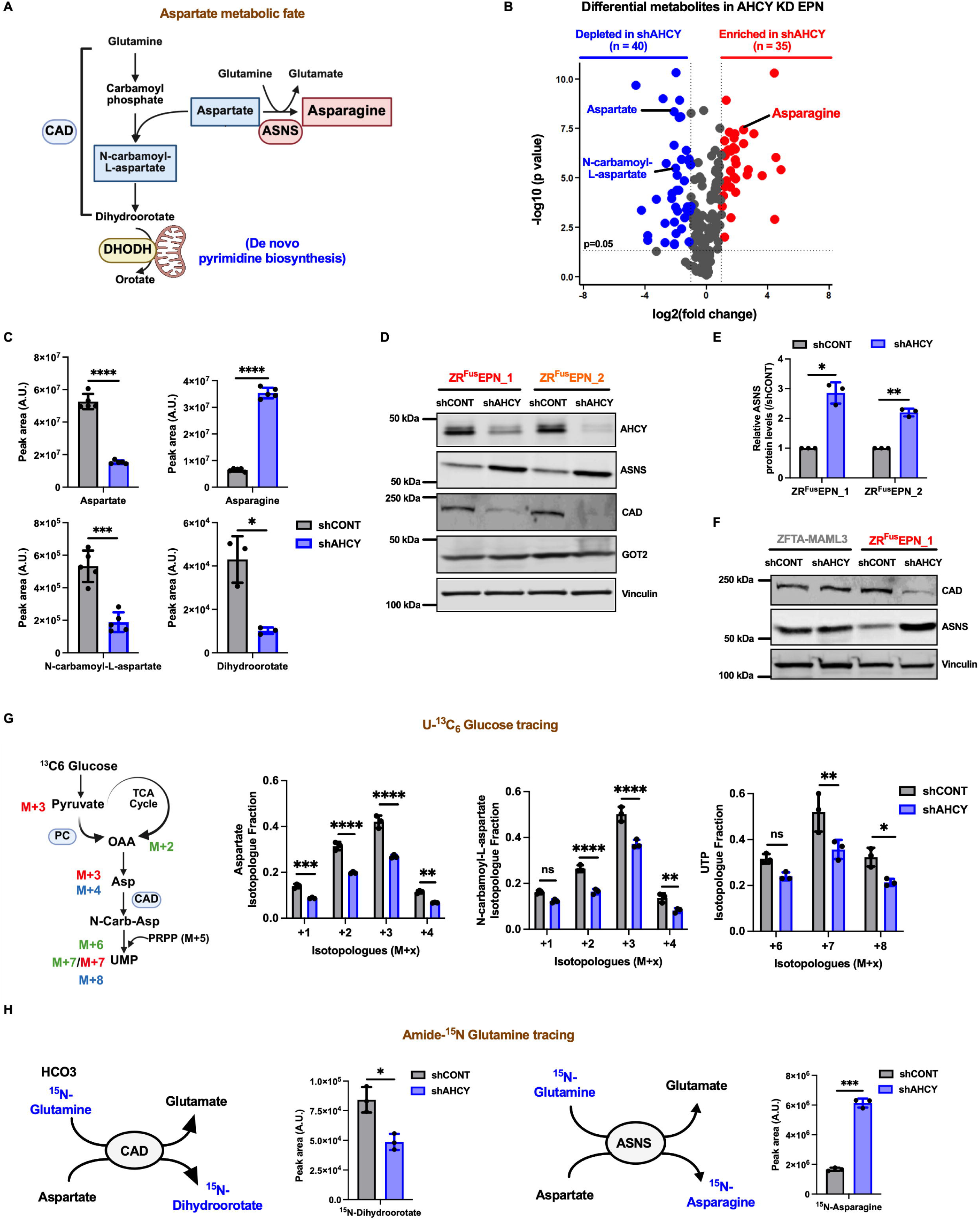
AHCY maintains aspartate utilization for de novo pyrimidine biosynthesis in ZFTA-RELA fusion tumors. (A) Schematic illustrating the metabolic fates of aspartate, including utilization for de novo pyrimidine biosynthesis through CAD-dependent N-carbamoyl-L-aspartate production and conversion to asparagine by asparagine synthetase (ASNS). CAD, carbamoyl-phosphate synthetase 2/aspartate transcarbamylase/dihydroorotase; DHODH, dihydroorotate dehydrogenase. (B) Volcano plot showing differential metabolites in patient-derived ZFTA-RELA fusion-positive (ZR^Fus^) supratentorial ependymoma (ST-EPN) cells following AHCY knockdown (KD) compared with control shRNA (shCONT). Significantly altered metabolites (p ≤ 0.05 and |log₂ fold change| ≥ 1) are highlighted in red (enriched) and blue (depleted). Aspartate, asparagine, and N-carbamoyl-L-aspartate are indicated. (C) Relative abundance of aspartate, asparagine, N-carbamoyl-L-aspartate, and dihydroorotate in patient-derived ZR^Fus^ cells expressing shCONT or shRNA targeting AHCY (shAHCY). (D) Representative immunoblots showing expression of AHCY, ASNS, CAD, and glutamic-oxaloacetic transaminase 2 (GOT2) in two independent patient-derived ZR^Fus^ cell lines expressing shCONT or shAHCY. Vinculin served as the loading control. (E) Densitometric quantification of ASNS protein expression following AHCY KD in ZR^Fus^ cell lines, normalized to shCONT. (F) Representative immunoblots showing CAD and ASNS expression following AHCY KD in patient-derived ZFTA-MAML3 fusion-positive and ZR^Fus^ ST-EPN cells. Vinculin served as the loading control. (G) Schematic of U-¹³C₆ glucose tracing into aspartate and downstream pyrimidine metabolites (left). Isotopologue distributions of ¹³C-labeled aspartate, N-carbamoyl-L-aspartate, and uridine triphosphate (UTP) in patient-derived ZR^Fus^ cells expressing shCONT or shAHCY (right). PC, pyruvate carboxylase; OAA, oxaloacetate; TCA, tricarboxylic acid; PRPP, phosphoribosyl pyrophosphate. (H) Schematic of amide-¹ N glutamine tracing into CAD-dependent pyrimidine biosynthesis and ASNS-mediated asparagine synthesis. Relative abundance of ¹ N-labeled dihydroorotate and ¹ N-labeled asparagine in patient-derived ZR^Fus^ cells expressing shCONT or shAHCY. For panel C, data are presented as mean ± SD from five independent samples per condition for aspartate, asparagine, and N-carbamoyl-L-aspartate and three independent samples per condition for dihydroorotate. For panels E, G, and H, data are presented as mean ± SD from three independent samples per condition. Statistical significance was determined using an unpaired two-tailed t-test with Welch’s correction for panels C, E, G, and H. *p < 0.05, **p < 0.01, ***p < 0.001, ****p < 0.0001; ns, not significant.

Functionally we assessed how AHCY regulates carbon flux through these competing pathways by performing stable isotope tracing using U-^13^C_6_-glucose in ZR^Fus^ cells expressing control or AHCY-targeting shRNA. In control cells, glucose-derived carbon was efficiently incorporated into aspartate, N-carbamoyl-L-aspartate, and downstream uridine triphosphate (UTP) isotopologues. In contrast, AHCY-depleted cells exhibited markedly reduced ¹³C incorporation into pyrimidine pathway metabolites, demonstrating impaired flux of glucose-derived carbon into de novo pyrimidine biosynthesis (**Fig. 4G**). Because both CAD-mediated pyrimidine synthesis and ASNS-mediated asparagine synthesis require glutamine-derived amide nitrogen, we next performed amide-¹ N-glutamine tracing. AHCY depletion significantly reduced ¹ N incorporation into dihydroorotate while simultaneously increasing the production of ¹ N-labeled asparagine (**Fig. 4H**), demonstrating a redistribution of glutamine-derived nitrogen away from pyrimidine synthesis and toward asparagine biosynthesis. Together, these carbon and nitrogen tracing studies establish that AHCY controls the metabolic fate of aspartate in ZR^Fus^ EPN.

### AHCY-mediated epigenetic regulation of asparagine metabolism

Given that AHCY promotes cellular methylation potential through the clearance of SAH^44^, we investigated whether AHCY regulates genes involved in de novo pyrimidine and asparagine synthesis in ZR^Fus^ EPN. A recent study in ZR tumor models demonstrate that H3K4me3 and H3K27me3 are enriched in ZR^Fus^ EPN tumors^18^. Similarly, immunoblot analysis of isogenic neural stem cell models revealed that maintenance of both histone marks is dependent on ZR fusion expression (**Fig. 5A**). We found that elevated AHCY expression is associated with increased H3K4me3 and H3K27me3, as AHCY loss resulted in a significant reduction in both modifications (**Fig. 5B**). Given the pronounced reduction in H3K4me3 and CAD expression upon AHCY depletion, we examined whether AHCY regulates CAD transcription through H3K4me3. ChIP-qPCR demonstrated significant H3K4me3 enrichment at the CAD promoter, which was markedly reduced following AHCY depletion (**Extended Data Fig. 4A**). This loss of promoter-associated H3K4me3 coincided with reduced CAD protein levels, consistent with impaired transcriptional activation (**Fig. 4D**).

**Fig. 5:**
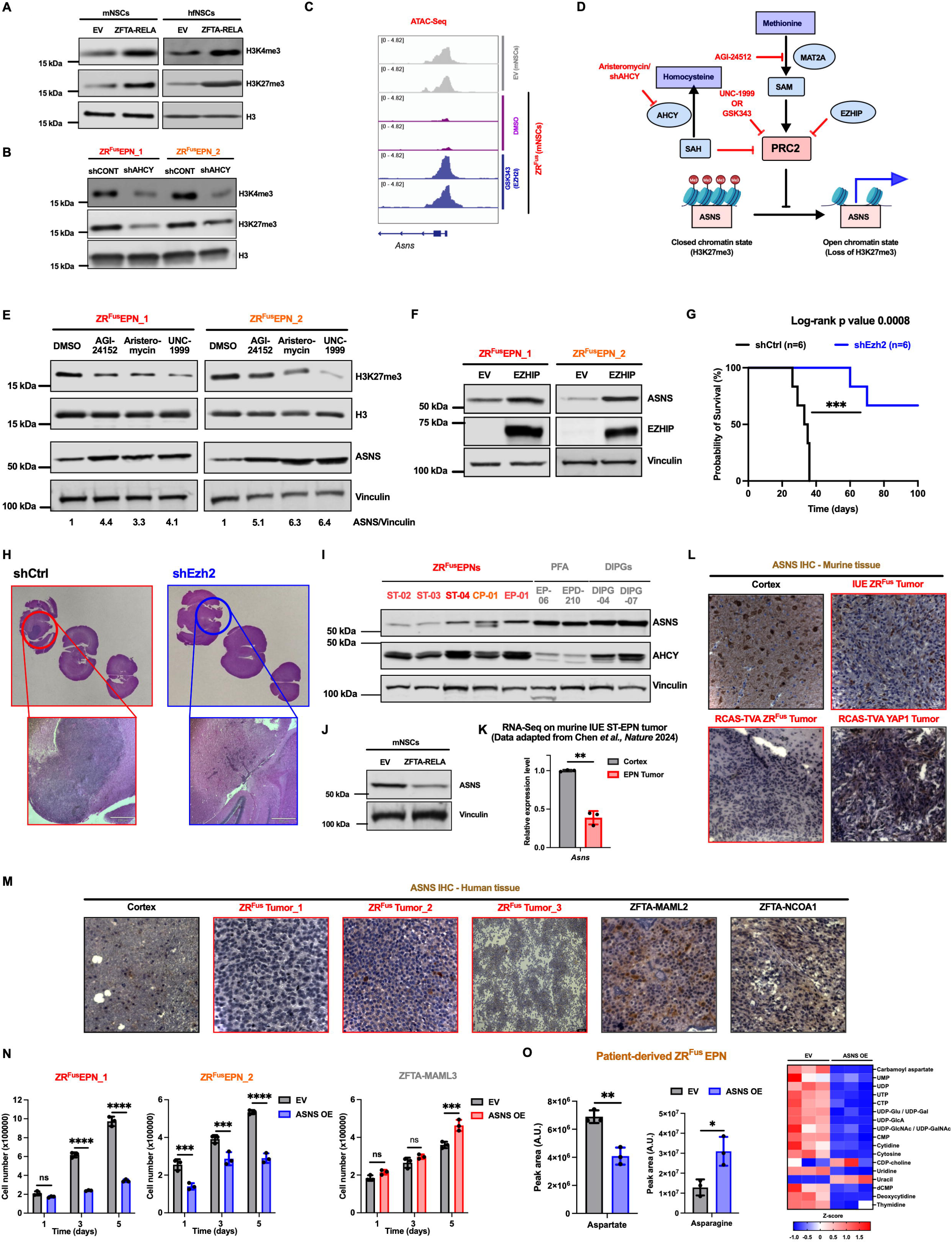
AHCY maintains H3K27me3-mediated repression of ASNS in ZFTA-RELA fusion ependymoma. (A) Representative immunoblots showing H3K4me3 and H3K27me3 levels in mouse neural stem cells (mNSCs) and human fetal neural stem cells (hfNSCs) expressing empty vector (EV) or ZFTA-RELA. Total H3 served as the loading control. (B) Representative immunoblots showing H3K4me3 and H3K27me3 levels in two independent patient-derived ZFTA-RELA fusion-positive (ZR^Fus^) supratentorial ependymoma (ST-EPN) cell lines expressing control shRNA (shCONT) or shRNA targeting AHCY (shAHCY). Total H3 served as the loading control. (C) Assay for transposase-accessible chromatin sequencing (ATAC-seq) tracks showing chromatin accessibility at the *Asns* locus in mNSCs expressing EV (top, light gray) or ZFTA-RELA following treatment with dimethyl sulfoxide (DMSO; middle, violet) or the EZH2 inhibitor GSK-343 (10 μM, 96 h; bottom, blue). (D) Schematic illustrating the proposed regulation of ASNS through the methionine–S-adenosylmethionine (SAM) axis and polycomb repressive complex 2 (PRC2). Pharmacological inhibition or depletion of MAT2A or AHCY, or inhibition of EZH2, is associated with loss of H3K27me3 and increased ASNS expression. MAT2A, methionine adenosyltransferase 2A; SAH, S-adenosylhomocysteine; EZHIP, EZH inhibitory protein. (E) Representative immunoblots showing H3K27me3 and ASNS levels in ZR^Fus^ cell lines following treatment with the MAT2A inhibitor AGI-24512 (5 μM), the AHCY inhibitor aristeromycin (2.5 μM), or the EZH1/2 inhibitor UNC-1999 (2 μM) for 72 h compared with DMSO control. Total H3 and vinculin served as loading controls. ASNS levels relative to vinculin are indicated. (F) Representative immunoblots showing ASNS expression following ectopic expression of EZHIP in two patient-derived ZR^Fus^ cell lines. Vinculin served as the loading control. (G) Kaplan–Meier survival analysis of immunocompetent C57BL/6J mice bearing orthotopic in utero electroporation (IUE)-derived murine ZR^Fus^ tumors expressing control shRNA (shCtrl) or shRNA targeting Ezh2 (shEzh2). (H) Representative hematoxylin and eosin (H&E)-stained coronal brain sections from mice bearing orthotopic shCtrl or shEzh2 ZR^Fus^ tumors. Insets show higher-magnification views of the tumor regions. Scale bar, 220 μm. (I) Representative immunoblots showing ASNS and AHCY expression across patient-derived ZR^Fus^ ST-EPN, posterior fossa A (PFA) ependymoma, and diffuse intrinsic pontine glioma (DIPG) cell lines. Vinculin served as the loading control. (J) Representative immunoblots showing ASNS expression in mNSCs expressing EV or ZFTA-RELA. Vinculin served as the loading control. (K) Relative expression of *Asns* in IUE-derived murine ZR^Fus^ tumors compared with normal cortex. Data were obtained from the RNA sequencing (RNA-seq) dataset reported by Chen *et al.* (*Nature*, 2024). (L) Representative immunohistochemistry (IHC) images showing ASNS expression in normal murine cortex, IUE-derived ZR^Fus^ tumors, RCAS-TVA-derived ZR^Fus^ tumors, and RCAS-TVA-derived YAP1 fusion-positive tumors. Scale bar, 50 μm. (M) Representative IHC images showing ASNS expression in normal human cortex and primary ST-EPN tumors harboring ZFTA-RELA, ZFTA-MAML2, or ZFTA-NCOA1 fusions. Scale bar, 50 μm, except ZR^Fus^ tumor 3, 100 μm. (N) Cell proliferation of two independent patient-derived ZR^Fus^ cell lines and a ZFTA-MAML3 fusion-positive ST-EPN cell line following ASNS overexpression (ASNS OE) compared with EV control. Cell number was measured over 5 days. (O) Relative abundance of aspartate and asparagine (left) and heatmap showing pyrimidine-associated metabolite abundance (right) in patient-derived ZR^Fus^ cells expressing EV or ASNS overexpression (ASNS OE). For panels K, N, and O, data are presented as mean ± SD from three independent samples per group or condition, respectively. Panel G includes six mice per group. Statistical significance was determined using an unpaired two-tailed t-test with Welch’s correction for panels K and O, two-way ANOVA with Šidák’s multiple-comparisons test for panel N, and the log-rank test for panel G. *p < 0.05, **p < 0.01, ***p < 0.001, ****p < 0.0001; ns, not significant.

We next sought to define the mechanism underlying AHCY-dependent repression of ASNS. Although amino acid starvation typically induces ASNS through GCN2-eIF2α-ATF4 signaling^45,46^ (**Extended Data Fig. 4B**), this pathway was not activated following AHCY depletion (**Extended Data Figs. 4C, D**), suggesting a non-canonical mechanism of ASNS regulation in ST-EPN. Given that H3K27me3 levels are enriched in ZR^Fus^ EPN^18^ and our finding indicated that AHCY is required for maintenance of increased H3K27me3 levels, we hypothesized that AHCY suppresses ASNS through PRC2-mediated histone methylation. Consistent with this, ATAC-seq analysis following EZH2 inhibition revealed increased chromatin accessibility at the ASNS locus, indicating H3K27me3-dependent repression (**Fig. 5C**). To further validate this regulatory axis, we independently perturbed multiple components governing H3K27me3 deposition, including MAT2A, which generates SAM for histone methylation; EZH2, the catalytic subunit of PRC2 responsible for H3K27me3 deposition; and AHCY, which hydrolyzes SAH and thereby sustains SAM-dependent methylation (**Fig. 5D**). Genetic or pharmacologic inhibition of these factors consistently induced ASNS expression (**Fig. 5E**). Furthermore, ectopic expression of EZHIP, a potent PRC2/EZH2 inhibitory protein, in ZR^Fus^ cells significantly increased ASNS expression (**Fig. 5F**). To further establish the specificity, we treated ZR^Fus^ cells with UNC0638, a selective inhibitor of the G9a/GLP H3K9 methyltransferase complex and found that inhibition of H3K9 methylation failed to induce ASNS expression (**Extended Data Fig. 4E**). Collectively, these independent validations reveal that ASNS repression in ST-EPN is mediated by AHCY and PRC2-dependent H3K27me3 in ST-EPN.

While low ASNS expression has previously been reported in acute lymphoblastic leukemia (ALL)^47,48^, inhibition of EZH2 failed to induce ASNS expression in ALL models (**Extended Data Fig. 4F**). These findings indicate that, although ASNS deficiency can occur across distinct malignancies, the underlying regulatory mechanisms are context dependent. In contrast to ALL, ASNS expression in ST-EPN appears to be uniquely governed by PRC2-mediated H3K27me3 repression. Collectively, our findings identify H3K27me3-dependent silencing of ASNS as a previously unrecognized, lineage-specific epigenetic mechanism in ZR fusion-driven ST-EPN.

Considering selective enrichment of H3K27me3 in ZR fusion-driven tumors and its critical role in ASNS suppression, we next evaluated the functional importance of chromatin modification H3K27me3 in vivo. To this end, we selectively knockdown EZH2 in an orthotopic ZR^Fus^ murine model and observed profound tumor reduction and significantly prolonged overall survival (**Figs. 5G, H**), demonstrating that H3K27me3-dependent chromatin regulation is required for tumor maintenance. Taken together, these data reveal that AHCY drives an integrated epigenetic and metabolic program in ZR^Fus^ EPN by linking methionine metabolism to H3K4me3-dependent activation of CAD and H3K27me3-mediated repression of ASNS.

### ZFTA-RELA EPN exhibit ASNS^low^ phenotype

We demonstrate that AHCY upregulation in ZFTA fusion-positive ependymoma epigenetically establishes an ASNS-low state. Indeed, patient-derived ZR^Fus^ models exhibited coordinated AHCY upregulation and ASNS suppression, a pattern not observed in other pediatric brain tumor controls (**Fig. 5I**). Moreover, expression of the ZR fusion in murine neural stem cells increased AHCY expression (**Fig. 3F**) while concomitantly reducing ASNS expression (**Fig. 5J**). This correlation was independently recapitulated in a published RNA-seq dataset from IUE-murine ZR^Fus^ ST-EPN model^37^ (GEO: GSE246033), which similarly showed elevated *Ahcy* (**Fig. 3G**) and reduced *Asns* expression (**Fig. 5K**). Collectively, these findings indicate that the ZFTA fusion oncogene drives establishment and maintenance of the ASNS-low phenotype. Consistently, immunohistochemical analysis of both murine and patient-derived ZR^Fus^ EPN tumors (**Figs. 5L, M**) revealed significantly reduced ASNS expression compared to other ST-EPN subtypes, including ZFTA-MAML2, ZFTA-NCOA1, and YAP1-MAMLD2, as well as in other pediatric brain tumor xenograft models such as medulloblastoma and diffuse intrinsic pontine glioma (DIPG) and normal cortex (**Figs. 5L, M and Extended Data Fig. 4G**). These findings indicate that low ASNS expression is a unique feature of ZR^Fus^ tumors.

To functionally evaluate necessity of ASNS^low^ status in ZR^Fus^ EPNs, we enforced ASNS expression in patient-derived ZR^Fus^ cells. Remarkably, ASNS overexpression profoundly inhibited proliferation of ZR^Fus^ EPN cells (**Fig. 5N**). In contrast, ASNS overexpression had no significant effect on the proliferation of ZFTA-MAML3 ST-EPN cells (**Fig. 5N**) or other pediatric brain tumor models, including pediatric high-grade glioma and DIPG (**Extended Data Fig. 4H**). Together, these findings identify ASNS suppression as a ZR fusion-specific metabolic vulnerability and suggest that maintenance of the ASNS-low state is required for the proliferative capacity of ZR fusion driven ependymoma. Next, we investigated the metabolic consequences of restoring ASNS expression in ZR^Fus^ ST-EPNs. Enforced ASNS expression resulted in depletion of intracellular aspartate and accumulation of asparagine (**Fig. 5O**), indicating increased flux through the asparagine biosynthetic pathway. Consistent with reduced aspartate availability, we observed marked depletion of pyrimidine biosynthetic intermediates and downstream nucleotides, including carbamoyl aspartate, UMP, UDP, UTP, and CTP (**Fig. 5O and Extended Data Fig. 4I and Supplementary Table 12**). Together, these data suggest that epigenetic suppression of ASNS preserves aspartate for de novo pyrimidine synthesis, whereas ASNS restoration redirects aspartate toward asparagine production at the expense of nucleotide biosynthesis.

### ZFTA-RELA EPN demonstrate extracellular asparagine dependency

Having established that ZR^Fus^ EPN maintain an ASNS-low state through AHCY-PRC2-H3K27me3-mediated repression (**Fig. 5**), we hypothesized that endogenous asparagine synthesis may be limited. To test this, we performed amide-¹ N-glutamine tracing in ST-EPN models and found significantly lower incorporation of glutamine-derived amide nitrogen into asparagine in ZR^Fus^ EPN cells compared with ZFTA-MAML3 EPN cells (**Fig. 6A**). As a result of reduced endogenous asparagine synthesis, asparagine may become a conditionally essential amino acid, rendering ZR^Fus^ EPN dependent on extracellular asparagine. We therefore examined extracellular asparagine uptake using U-¹³C₄-asparagine and found ZR^Fus^ EPN cells exhibited significantly greater intracellular accumulation of ¹³C₄-asparagine, accompanied by a corresponding reduction of labeled asparagine remaining in the culture medium, compared with ZFTA–MAML3 cells (**Fig. 6B**). Consistent with this increased uptake, increasing concentration of exogenous asparagine significantly enhanced proliferation of ZR^Fus^ EPN models, whereas asparagine addition had no significant effect on proliferation of ZFTA-MAML3 EPN or other pediatric brain tumor models (**Fig. 6C and Extended Data Fig. 5A**). These data support ZR^Fus^ EPN specific dependence on exogenous asparagine to sustain growth.

**Fig. 6:**
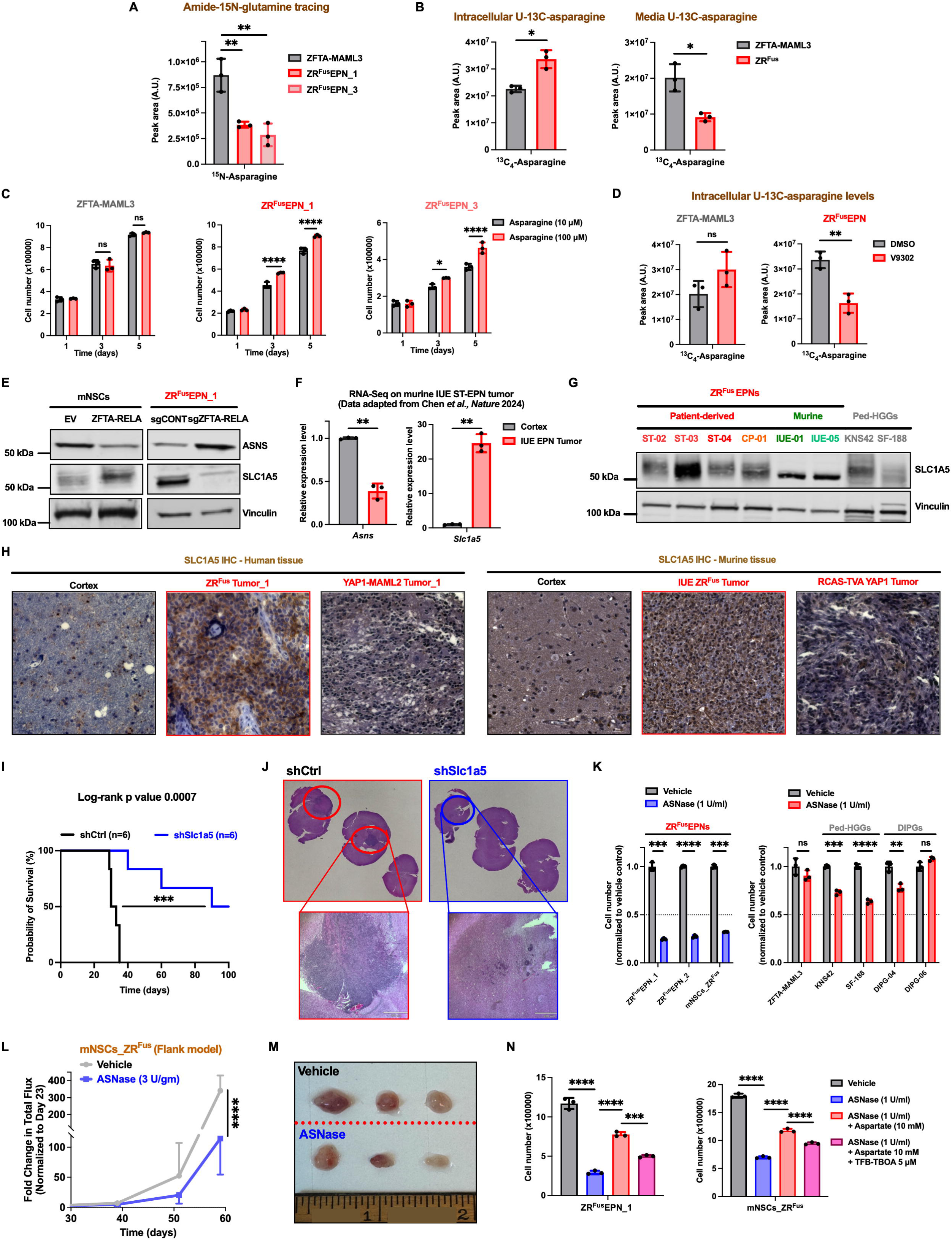
ZFTA-RELA fusion ependymoma depends on SLC1A5-mediated asparagine uptake. (A) Amide-¹ N-glutamine tracing showing ¹ N incorporation into asparagine in patient-derived ZFTA-RELA fusion-positive (ZR^Fus^) supratentorial ependymoma (ST-EPN) cells compared with ZFTA-MAML3 fusion-positive ST-EPN cells. (B) Intracellular and extracellular abundance of U-¹³C-asparagine following supplementation with U-¹³C-asparagine in patient-derived ZR^Fus^ cells compared with ZFTA-MAML3 fusion-positive ST-EPN cells. (C) Cell proliferation of two independent patient-derived ZR^Fus^ cell lines and a ZFTA-MAML3 fusion-positive ST-EPN cell line cultured with 10 μM or 100 μM asparagine. Cell number was measured over 5 days. (D) Intracellular U-¹³C-asparagine abundance in patient-derived ZR^Fus^ and ZFTA-MAML3 fusion-positive ST-EPN cells following 8 h treatment with dimethyl sulfoxide (DMSO) or the SLC1A5 inhibitor V-9302. (E) Representative immunoblots showing ASNS and SLC1A5 expression in mouse neural stem cells (mNSCs) expressing empty vector (EV) or ZFTA-RELA and in patient-derived ZR^Fus^ cells expressing control single-guide RNA (sgRNA; sgCONT) or sgRNA targeting ZFTA-RELA (sgZFTA-RELA). Vinculin served as the loading control. (F) Relative expression of *Asns* and *Slc1a5* in IUE-derived murine ZR^Fus^ tumors compared with normal cortex. Data were obtained from the RNA sequencing (RNA-seq) dataset reported by Chen *et al.* (Nature, 2024). (G) Representative immunoblots showing SLC1A5 expression across patient-derived and IUE-derived murine ZR^Fus^ cell lines compared with pediatric high-grade glioma (Ped-HGG) cell lines. Vinculin served as the loading control. (H) Representative immunohistochemistry (IHC) images showing SLC1A5 expression in normal human cortex, primary ZR^Fus^ and YAP1-MAML2 fusion-positive ST-EPN tumors (left), and normal murine cortex, in utero electroporation (IUE)-derived ZR^Fus^ tumors, and RCAS-TVA-derived YAP1 fusion-positive tumors (right). Scale bars, 50 μm. (I) Kaplan-Meier survival analysis of immunocompetent C57BL/6J mice bearing orthotopic IUE-derived murine ZR^Fus^ tumors expressing control shRNA (shCtrl) or shRNA targeting Slc1a5 (shSlc1a5). (J) Representative hematoxylin and eosin (H&E)-stained coronal brain sections from mice bearing orthotopic shCtrl or shSlc1a5 ZR^Fus^ tumors. Insets show higher-magnification views of the tumor regions. Scale bar, 220 μm. (K) Relative cell number of two patient-derived ZR^Fus^ cell lines and ZFTA-RELA-expressing mNSCs following treatment with vehicle or L-asparaginase (ASNase; 1 U ml⁻¹) for 96 h (left). Relative cell number of a ZFTA-MAML3 fusion-positive ST-EPN cell line, Ped-HGG cell lines, and diffuse intrinsic pontine glioma (DIPG) cell lines following treatment under the same conditions (right). Values were normalized to the corresponding vehicle control. (L) Bioluminescence-based growth of subcutaneous tumors generated from ZFTA-RELA-expressing mNSCs in mice treated with vehicle or ASNase (3 U g⁻¹) every other day, three times per week. Tumor burden was monitored longitudinally by total flux and normalized to day 23. (M) Representative images of excised subcutaneous tumors generated from ZFTA-RELA-expressing mNSCs in mice treated with vehicle or ASNase and collected on day 60. (N) Cell number of patient-derived ZR^Fus^ cells and ZFTA-RELA-expressing mNSCs following 96 h treatment with vehicle, ASNase (1 U ml⁻¹), or ASNase supplemented with aspartate (10 mM), or ASNase plus aspartate in the presence of the aspartate transport inhibitor TFB-TBOA (5 μM). For panels A–D, F, K, and N data are presented as mean ± SD from three independent samples per group or condition. For panel L, data are presented as mean ± SD from three mice per group. Panel I includes six mice per group. Unless otherwise indicated, all in vitro ASNase experiments were performed in Neurobasal medium supplemented with asparagine (0.1 mM). For 96 h ASNase treatments, fresh vehicle or ASNase was added after 48 h. Statistical significance was determined using one-way ANOVA with Dunnett’s multiple-comparisons test for panels A and N, an unpaired two-tailed t-test with Welch’s correction for panels B, D, F, and K, two-way ANOVA with Šidák’s multiple-comparisons test for panels C and L, one-way ANOVA with Šidák’s multiple-comparisons test for panel N, and the log-rank test for panel I. *p < 0.05, **p < 0.01, ***p < 0.001, ****p < 0.0001; ns, not significant.

We then investigated the mechanism by which ZR^Fus^ EPN cells acquire extracellular asparagine. A recent study of ST-EPN identified SLC1A5 as a key transporter that is upregulated to support production of the oncometabolite itaconate^18^. SLC1A5 is a solute carrier protein that mediates transport of neutral amino acids, including glutamine and asparagine^49–51^, and its overexpression has been implicated in maintaining intracellular amino acid homeostasis in malignant cells^52,53^. Although SLC1A5 transports both glutamine and asparagine, it has been studied primarily as a glutamine transporter because glutamine is widely regarded as a conditionally essential amino acid for proliferating cancer cells. In contrast, asparagine is generally considered nonessential, as its intracellular levels are maintained through ASNS-mediated de novo synthesis in most cancer models. Notable exceptions include acute lymphoblastic leukemia (ALL) and ZR fusion-positive ST-EPNs as demonstrated here, where ASNS expression is suppressed. These observations suggest that the functional importance of SLC1A5 may extend beyond glutamine transport in ASNS^low^ tumors, where it may also contribute to maintaining intracellular asparagine homeostasis. To determine whether SLC1A5 mediates asparagine uptake in ZR^Fus^ EPN cells, we supplemented culture media with exogenous U-^13^C_4_-asparagine and quantified intracellular labeled asparagine levels in the presence of vehicle control or V9302, a selective SLC1A5 inhibitor. LC-MS/MS analysis demonstrated that intracellular U-^13^C_4_-asparagine levels were significantly reduced following V9302 treatment in ZR^Fus^ EPN cells compared with ZFTA–MAML3 EPN cells (**Fig. 6D**), indicating that SLC1A5 is a major contributor to extracellular asparagine uptake.

To define the relationship between ASNS and SLC1A5 expression in ZR fusion-positive ependymoma, we examined complementary isogenic models. Ectopic expression of the ZR fusion in neural stem cells induced SLC1A5 expression while suppressing ASNS, whereas knockout of ZR fusion oncogene in patient-derived models resulting in reduced SLC1A5 and increased ASNS expression (**Fig. 6E**). These findings indicate that the ZR fusion establishes and maintains a transcriptional state characterized by high SLC1A5 and low ASNS expression. Consistent with this model, analysis of a recently published RNA-seq dataset from an IUE-derived ZR fusion-positive ST-EPN model revealed markedly elevated *Slc1a5* and reduced *Asns* expression in tumors relative to normal cortex^37^ (GEO: GSE246033) (**Fig. 6F**). Importantly, single-cell RNA sequencing analyses from both human and murine ST-EPN tumors demonstrated that malignant tumor cell populations were characterized by high SLC1A5 and low ASNS expression (**Extended Data Figs. 5B, C**).

SLC1A5 expression was also elevated in patient-derived and murine ZR^Fus^ EPN models compared with other brain tumor models (**Fig. 6G**). Consistently, patient and murine ZR^Fus^ ST-EPN tumors exhibited increased SLC1A5 expression, whereas expression was comparatively low in normal cortex and YAP1 fusion-positive ST-EPN tumors (**Fig. 6H**). We next sought to determine how SLC1A5 expression is regulated in ST-EPN. To this end, we interrogated previously published ZR fusion ChIP-seq datasets derived from murine ST-EPN model^27^. We found that the upstream regulatory region of SLC1A5 is directly bound by the ZR fusion oncoprotein (**Extended Data Fig. 5D**), suggesting transcriptional activation by ZR fusion. Moreover, H3K27ac ChIP-seq data from patient-derived ST-EPN samples revealed a hyperacetylated super-enhancer-like region upstream of SLC1A5^7^ (**Extended Data Fig. 5E**), further supporting active transcriptional regulation of SLC1A5.

To validate SLC1A5 dependency in tumor maintenance in vivo, we orthotopically implanted murine ZR^Fus^ EPN cells expressing either control shRNA or shRNA targeting Slc1a5 into the cerebral cortex of immunocompetent mice. Depletion of Slc1a5 significantly reduced tumor burden and prolonged survival compared with control tumors (**Figs. 6I, J**), demonstrating that SLC1A5 is functionally required for ZR^Fus^ EPN growth in vivo. Together, these findings suggest that SLC1A5 plays a critical role in mediating asparagine uptake and supports the proliferation of ZR fusion driven ST-EPN cells through regulation of asparagine homeostasis.

### ZFTA-RELA EPN is sensitive to L-asparaginase

L-asparaginase (ASNase), an FDA-approved enzyme therapy that depletes extracellular L-asparagine, is a frontline treatment for acute lymphoblastic leukemia^48,54^. Given that asparagine is conditionally required for ST-EPN proliferation, we investigated whether ASNase could be therapeutically exploited in ST-EPN tumors. ASNase markedly reduced the proliferation of patient-derived ZR^Fus^ EPN and ZR-expressing mNSCs, whereas other brain tumor cells displayed little or no sensitivity (**Fig. 6K and Extended Data Fig. 5F**). Because the currently available pegylated form of ASNase is not blood-brain barrier (BBB) permeable^55^, we evaluated its antitumor activity using a subcutaneous ZR tumor model. ASNase treatment significantly suppressed tumor growth and reduced tumor size compared with vehicle-treated controls (**Figs. 6L, M**), suggesting that brain-penetrant or CNS-targeted ASNase delivery strategies may represent a potential therapeutic approach for ZR fusion-positive ST-EPN tumors.

Extracellular asparagine depletion by ASNase treatment induced robust upregulation of endogenous ASNS expression (**Extended Data Fig. 5G**). Although untreated ZR fusion-positive EPN cells were largely insensitive to the ASNS inhibitor ASX-173, ASNase pretreatment markedly increased sensitivity, indicating that endogenous ASNS becomes a compensatory dependency when extracellular asparagine is limited (**Extended Data Fig. 5H**). Although sustained ASNS overexpression impaired ZR^Fus^ EPN proliferation, the acute induction of ASNS following extracellular asparagine depletion may represent an adaptive response that helps maintain intracellular asparagine availability during nutrient stress.

Given that ASNS-mediated asparagine synthesis consumes aspartate as a key substrate, we hypothesized that aspartate availability may become limiting following ASNase treatment and the consequent reactivation of endogenous ASNS. To test this, we supplemented ASNase-treated ST-EPN cells with a supraphysiological concentration of aspartate (10 mM), which significantly rescued the ASNase-induced proliferation defect (**Fig. 6N and Extended Data Fig. 5I**). Notably, inhibition of aspartate transport with TFB-TBOA impaired this rescue, demonstrating that uptake of extracellular aspartate is necessary to support proliferation under these conditions (**Fig. 6N**). These findings are consistent with a model in which repression of ASNS preserves intracellular aspartate pools for de novo pyrimidine synthesis enabling nucleotide production and sustained proliferation in ZR fusion-positive ST-EPN.

### De novo pyrimidine biosynthesis is an actionable vulnerability in ZFTA-RELA EPN

Because CAD, the initiating enzyme complex of de novo pyrimidine biosynthesis, and ASNS compete for the same metabolic substrates, aspartate and glutamine, we hypothesized that ASNS repression may promote pyrimidine production by increasing substrate availability for CAD. Consistent with this hypothesis, enforced ASNS expression in ZR fusion-positive EPN cells resulted in broad depletion of pyrimidine pathway intermediates and nucleotides (**Fig. 5O**), indicating that diversion of aspartate toward asparagine synthesis constrains de novo pyrimidine biosynthesis.

Metabolomic pathway analysis identified de novo pyrimidine biosynthesis as the most significantly enriched metabolic pathway in murine ZR^Fus^ EPN tumors (**Fig. 7A**). At the metabolite level, the pyrimidine biosynthetic intermediates N-carbamoyl-L-aspartate and dihydroorotate were elevated in murine ZR^Fus^ tumors relative to normal cortex and were correspondingly depleted following ZR fusion knockout in patient models (**Fig. 7B and Extended Data Figs. 3I, J**). We next asked whether aspartate is preferentially directed toward de novo pyrimidine biosynthesis in ZR^Fus^ cells. To this end, we performed U-^13^C_6_-glucose tracing to examine the incorporation of glucose-derived carbon into aspartate, pyrimidine intermediates, and UMP. Glucose-derived M+4 aspartate generates M+4 N-carbamoyl-L-aspartate and, following loss of one aspartate-derived carbon and addition of M+5 ribose from PRPP, gives rise to UMP M+8. We therefore used the measured aspartate isotopologue distribution to predict UMP labeling under a diffuse metabolic model in which pathway intermediates freely equilibrate with the bulk intracellular metabolite pool. Under this model, the abundance of UMP M+8 should reflect the relative abundance of M+4 aspartate available within the bulk intracellular pool^56^ (**Fig. 7C**). Consistent with the initial incorporation of aspartate into the pathway, the relative M+1 to M+4 isotopologue distributions were similar between aspartate and N-carbamoyl-L-aspartate (**Extended Data Fig. 6A**). However, comparison of the predicted and experimentally observed UMP isotopologue distributions revealed a marked enrichment of UMP M+8 above that predicted by the diffuse model (**Fig. 7D**). This finding support preferential channeling of aspartate into de novo pyrimidine biosynthesis, potentially facilitated by spatial organization of pathway enzymes within a pyrimidinosome complex^57^ (**Fig. 7E**).

**Fig. 7:**
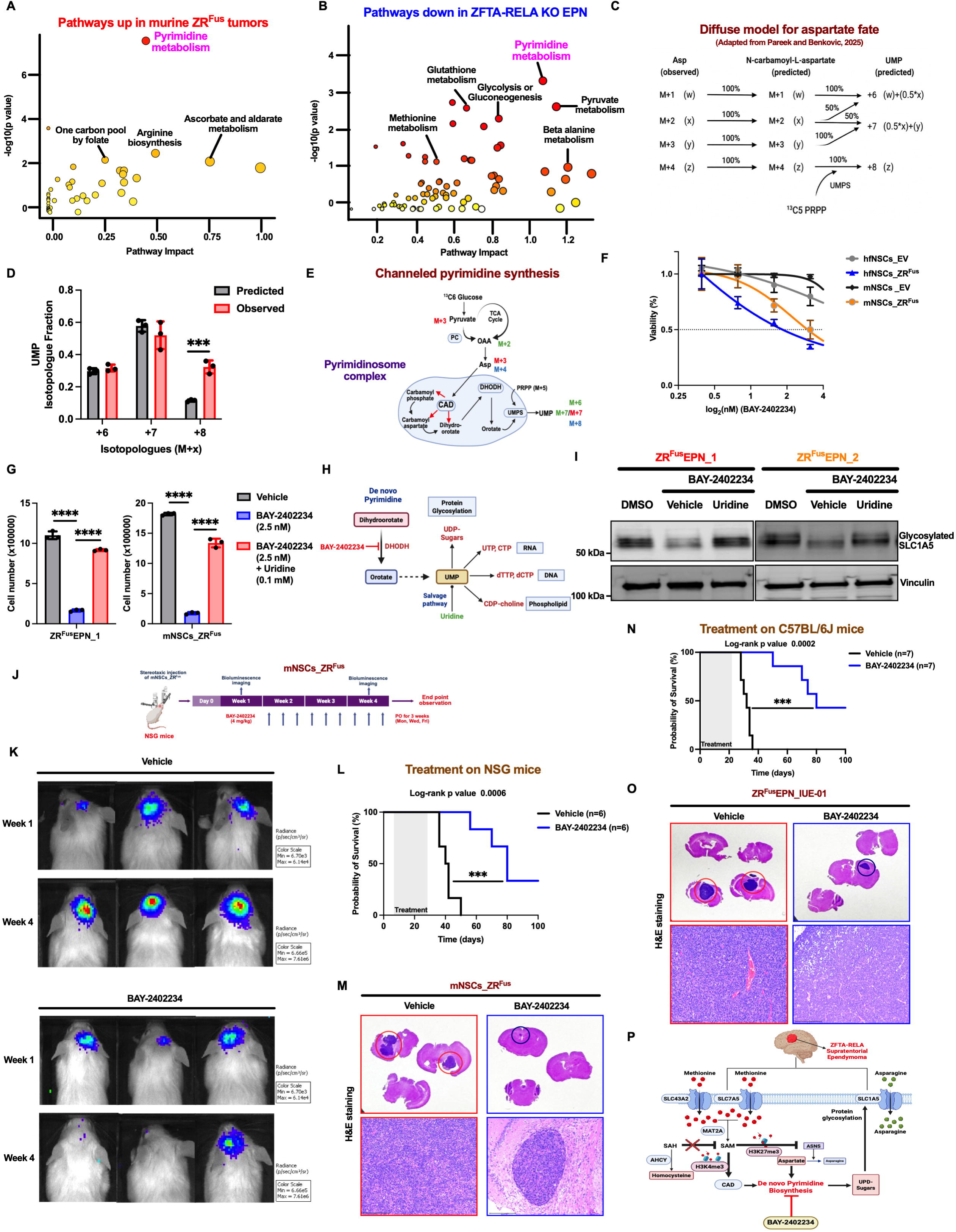
De novo pyrimidine biosynthesis is a therapeutic vulnerability in ZFTA-RELA fusion ependymoma. (A) Metabolic pathway enrichment analysis of metabolites significantly enriched in murine ZFTA-RELA fusion-positive (ZR^Fus^) tumors compared with normal cortex. Pyrimidine metabolism was the most significantly enriched pathway. (B) Integrated metabolomic and transcriptomic pathway analysis following ZFTA-RELA knockout in patient-derived ZR^Fus^ cells, incorporating significantly depleted metabolites (p ≤ 0.05 and log₂ fold change ≤ −0.58) and downregulated genes (false discovery rate (FDR) ≤ 0.05 and log₂ fold change ≤ −1). Pyrimidine metabolism was the most significantly suppressed pathway. (C) Schematic illustrating the predicted isotopologue distribution of uridine monophosphate (UMP) generated from U-¹³C₆-glucose-derived aspartate and ¹³C₅-labeled phosphoribosyl pyrophosphate (PRPP) under a diffuse model of aspartate utilization. Adapted from Pareek and Benkovic (*Nature Communications*, 2025). (D) Comparison of predicted and experimentally observed +6, +7, and +8 UMP isotopologue fractions following U-¹³C₆-glucose tracing in patient-derived ZR^Fus^ cells. (E) Schematic illustrating the proposed channeling of glucose-derived aspartate into de novo pyrimidine biosynthesis through a proposed pyrimidinosome complex. PC, pyruvate carboxylase; OAA, oxaloacetate; Asp, aspartate; TCA, tricarboxylic acid; CAD, carbamoyl-phosphate synthetase 2/aspartate transcarbamylase/dihydroorotase; DHODH, dihydroorotate dehydrogenase; UMPS, uridine monophosphate synthase; PRPP, phosphoribosyl pyrophosphate. (F) Dose-response curves showing viability of mouse neural stem cells (mNSCs) and human fetal neural stem cells (hfNSCs) expressing empty vector (EV) or ZFTA-RELA following 72 h treatment with increasing concentrations of the DHODH inhibitor BAY-2402234. (G) Cell number of patient-derived ZR^Fus^ cells (left) and ZFTA-RELA-expressing mNSCs (right) following treatment with vehicle, BAY-2402234 (2.5 nM), or BAY-2402234 plus uridine (0.1 mM). (H) Schematic illustrating DHODH-dependent de novo pyrimidine biosynthesis and downstream utilization of pyrimidine nucleotides for RNA and DNA synthesis, UDP-sugar production and protein glycosylation, and phospholipid synthesis, together with the uridine salvage pathway. (I) Representative immunoblots showing glycosylated SLC1A5 expression in two patient-derived ZR^Fus^ cell lines following treatment with DMSO, BAY-2402234, or BAY-2402234 plus uridine. Vinculin served as the loading control. (J) Schematic of the orthotopic tumor model generated from ZFTA-RELA-expressing mNSCs and the BAY-2402234 treatment schedule in NOD scid gamma (NSG) mice. Mice received BAY-2402234 (4 mg/kg) by oral gavage three times per week (Monday, Wednesday, and Friday) for 3 weeks. (K) Representative bioluminescence images of mice bearing orthotopic tumors generated from ZFTA-RELA-expressing mNSCs and treated with vehicle or BAY-2402234 at weeks 1 and 4. (L) Kaplan-Meier survival analysis of NSG mice bearing orthotopic tumors generated from ZFTA-RELA-expressing mNSCs and treated with vehicle or BAY-2402234 (n = 6 per group). (M) Representative hematoxylin and eosin (H&E)-stained coronal brain sections from mice bearing orthotopic tumors generated from ZFTA-RELA-expressing mNSCs and treated with vehicle or BAY-2402234. Boxed regions are shown at higher magnification. (N) Kaplan-Meier survival analysis of immunocompetent C57BL/6J mice bearing orthotopic IUE-derived ZR^Fus^ tumors treated with vehicle or BAY-2402234 (n = 7 per group). (O) Representative H&E-stained coronal brain sections from mice bearing orthotopic IUE-derived ZR^Fus^ tumors treated with vehicle or BAY-2402234. Boxed regions are shown at higher magnification. (P) Schematic summarizing the metabolic program in ZR^Fus^ supratentorial ependymoma (ST-EPN), linking enhanced methionine metabolism with H3K27me3-mediated ASNS repression, H3K4me3-associated regulation of CAD, SLC1A5-mediated asparagine uptake, aspartate utilization for de novo pyrimidine biosynthesis, and downstream UDP-sugar production. For panels D, F, and G, data are presented as mean ± SD from three independent samples per condition. For panel A, metabolomic analysis was performed using four biological replicates per group. For panel B, metabolomic and transcriptomic analyses were performed using three independent samples per group. Panels L and N include six and seven mice per group, respectively. Statistical significance was determined using an unpaired two-tailed t-test with Welch’s correction for panel D, one-way ANOVA with Šidák’s multiple-comparisons test for panel G, and the log-rank test for panels L and N. ***p < 0.001, ****p < 0.0001.

These findings suggest that enhanced de novo pyrimidine biosynthesis represents a metabolic vulnerability and actionable dependency in ZR^Fus^ EPN. We evaluated the sensitivity of ZR^Fus^ cells to inhibition of dihydroorotate dehydrogenase (DHODH), rate limiting enzyme in de novo pyrimidine synthesis (**Extended Data Fig. 6B**), that has been therapeutically targeted in several cancers, including brain tumors^58–61^. Patient-derived ZR^Fus^ models exhibited nanomolar level sensitivity to DHODH inhibition by BAY-2402234 (IC50) and ectopic expression of ZR expression in both hfNSCs and mNSCs also showed markedly increased sensitivity to BAY-2402234 relative to their respective controls (**Fig. 7F and Extended Data Fig. 6C**). Importantly, exogenous supplementation of uridine rescued BAY-2402234-induced growth inhibition in both patient-derived ZR^Fus^ cells and ZR-expressing mNSCs (**Fig. 7G and Extended Data Fig. 6D**), demonstrating that the antiproliferative effects of DHODH inhibition result predominantly from pyrimidine nucleotide depletion.

Beyond supplying precursors for DNA and RNA synthesis, de novo pyrimidine biosynthesis generates UDP-sugars required for protein glycosylation, which regulates protein folding, stability, trafficking, and cell-surface expression^62,63^ (**Fig. 7H**). N-glycosylation is particularly important for SLC1A5 maturation and function^64^. Given that SLC1A5 is upregulated and required for tumor growth in ZR^Fus^ ST-EPN (**Figs. 6G, H**), we asked whether pyrimidine metabolism maintains SLC1A5 glycosylation. BAY-2402234 reduced glycosylated SLC1A5, which was restored by uridine supplementation (**Fig. 7I**). Similarly, enforced ASNS expression depleted UDP-sugars, including UDP-GlcNAc, and reduced SLC1A5 glycosylation (**Fig. 5O and Extended Data Fig. 6E**). Together, these findings suggest that de novo pyrimidine biosynthesis supports both nucleotide production and UDP-sugar availability, thereby maintaining SLC1A5 glycosylation and asparagine uptake in ZR fusion-driven ependymoma.

We next evaluated the therapeutic efficacy of DHODH inhibition in two independent in vivo models of ZR^Fus^ EPN. First, mNSCs expressing ZR were orthotopically engrafted into NSG mice, which were subsequently randomized to receive either vehicle or BAY-2402234, a blood–brain barrier-penetrant DHODH inhibitor. Animals were treated with BAY-2402234 (4 mg kg⁻¹, oral gavage) for three weeks, and tumor progression was monitored by longitudinal bioluminescence imaging (**Figs. 7J, K**). BAY-2402234 treatment significantly suppressed tumor growth and prolonged survival compared with vehicle-treated controls (**Fig. 7L**). Histological analyses further demonstrated a marked reduction in tumor burden in treated animals (**Fig. 7M and Extended Data Fig. 6F**). To independently validate the therapeutic potential of DHODH inhibition in an immunocompetent setting, we utilized a syngeneic ZR^Fus^ model in C57BL/6J mice. Following orthotopic implantation of IUE-derived ZR^Fus^ tumor cells, tumor-bearing animals were randomized to receive vehicle or BAY-2402234 treatment. Consistent with results obtained in NSG mice, DHODH inhibition significantly prolonged survival in immunocompetent animals relative to vehicle-treated controls (**Fig. 7N**). Histological analyses confirmed substantially reduced tumor burden in BAY-2402234-treated mice (**Fig. 7O and Extended Data Fig. 6F**).

Collectively, these findings establish de novo pyrimidine biosynthesis as a critical metabolic vulnerability and actionable target in ZR fusion-positive EPN (**Fig. 7P**). The brain-penetrant DHODH inhibitor BAY-2402234 robustly reduced tumor burden and prolonged survival across multiple preclinical models, providing a compelling foundation for DHODH inhibitor clinical translation in ZFTA-RELA fusion-positive EPN.

## Discussion

Pediatric brain tumors are increasingly recognized as diseases in which developmental cell states, epigenetic regulation, and metabolism converge to drive malignant transformation^17,18,21^. Ependymomas are central nervous system malignancies that arise predominantly in children and adolescents and are characterized by profound epigenetic dysregulation and recurrent oncogenic fusion proteins^4,5,27^. Although oncogenic fusion proteins define the molecular identity of supratentorial ependymomas (ST-EPNs), how they reprogram cellular metabolism to sustain tumor growth remains incompletely understood. Here, through integrated metabolomic, transcriptomic, and functional analyses of patient tumors and genetically engineered models, we identify a previously unrecognized metabolic–epigenetic program through which ZFTA–RELA (ZR) ST-EPN rewires methionine metabolism to regulate chromatin state, amino acid uptake, and nucleotide biosynthesis. Dysregulated methionine metabolism has been reported across multiple cancer types^19,29,34^; but the mechanisms by which ZR oncogene establish and maintain methionine dependence remain poorly understood.

We demonstrate that ZR enhances methionine uptake through SLC7A5 and regulates methionine-cycle activity by increasing AHCY expression. We identify AHCY as a critical driver linking methionine-cycle activity to the transcriptional and metabolic programs required for tumor maintenance. Although methionine metabolism is considered critical for one-carbon metabolism and serves as a source of methyl donors for chromatin regualtion^29,30^, our data demonstrate that maintaining methylation potential through AHCY-mediated S-adenosylhomocysteine clearance is also critical and establish AHCY as a regulator of transcriptional and downstream metabolic reprogramming in ZR ST-EPN. A recent study demonstrated that itaconate stabilizes H3K4me3 in ZR ST-EPN through inhibition of histone demethylases^18^. Our findings extend this model by showing that enhanced methionine uptake and AHCY activity contribute to the maintenance of both H3K4me3 and H3K27me3. We further demonstrate that this epigenetic state drives a distinct metabolic program characterized by PRC2-dependent ASNS repression, altered aspartate flux, and enhanced de novo pyrimidine biosynthesis.

Furthermore, our results establish the ASNS-low state as a defining metabolic feature of ZR ST-EPN. Many cancers retain or increase ASNS-mediated asparagine synthesis. In glioblastoma, for example, ASNS is frequently amplified, and elevated ASNS expression is associated with poor patient survival^65^. The ASNS-low phenotype of ZR ST-EPN is particularly compelling because ASNS is normally highly expressed in the human brain^66^, and loss-of-function mutations in *ASNS* cause asparagine synthetase deficiency, a severe neurodevelopmental disorder characterized by congenital microcephaly^67^. Enforced ASNS expression selectively impaired the proliferation and metabolic fitness of ZR ST-EPN cells, demonstrating that maintenance of ASNS-low state is functionally important for supporting the anabolic demands of tumor growth. Although reduced ASNS expression has also been observed in a subset of acute lymphoblastic leukemia (ALL), we did not observe H3K27me3-dependent regulation of ASNS in ALL models, indicating that the ASNS-low state in ZR ST-EPN is established through a distinct epigenetic mechanism.

A direct consequence of ASNS downregulation is that ZR tumors become conditionally dependent on extracellular asparagine availability. We demonstrate that these tumors compensate for limited endogenous asparagine synthesis by increasing SLC1A5-mediated amino acid uptake, thereby establishing an ASNS^low^/SLC1A5^high^ metabolic state. This adaptation creates a therapeutically exploitable vulnerability. Genetic or pharmacological disruption of SLC1A5-dependent asparagine acquisition impaired tumor growth, while depletion of extracellular asparagine with L-asparaginase displayed marked antitumor activity. Identification of this previously unrecognized asparagine dependency is particularly noteworthy given the long-standing clinical success of asparagine-depletion strategies in acute lymphoblastic leukemia^54^. However, the limited central nervous system penetration of currently available L-asparaginase formulations underscores the need to develop brain-penetrant or brain-directed strategies that exploit this dependency in ZR ST-EPN.

Our metabolic analyses identified de novo pyrimidine biosynthesis as a critical metabolic dependency in ZR ST-EPN. Beyond supplying nucleotides for DNA and RNA synthesis, pyrimidine-derived metabolites support phospholipid biosynthesis and protein glycosylation, thereby supporting multiple processes required for tumor proliferation. ASNS suppression thus provides selective advantage by limiting the consumption of aspartate and glutamine for endogenous asparagine synthesis and redirecting these metabolites for pyrimidine biosynthesis. This metabolic regulation further compensates for reduced endogenous asparagine synthesis by upregulating SLC1A5 and thereby enhancing extracellular glutamine and asparagine uptake. Given that N-glycosylation is required for proper SLC1A5 maturation and transport activity, increased pyrimidine flux can also support amino acid homeostasis through nucleotide-sugar production and maintenance of SLC1A5 glycosylation. Thus, ASNS suppression may enable ZR ST-EPN cells to redirect metabolic resources toward pyrimidine biosynthesis, coordinating nucleotide production, protein glycosylation, and extracellular asparagine acquisition to support tumor proliferation. Consistent with this model, pharmacological inhibition of DHODH produced robust antitumor activity across multiple preclinical models, providing a strong rationale for therapeutically targeting de novo pyrimidine biosynthesis dependency in ZR ependymoma.

In summary, we demonstrate that the ZR fusion establishes an integrated metabolic–epigenetic program that couples methionine uptake and AHCY-dependent metabolic reprograming of pyrimidine biosynthesis and asparagine synthesis. More broadly, our findings reveal how an oncogenic fusion protein can exploit metabolic pathways to reshape chromatin regulation and reprogram downstream metabolic pathways that fuel aggressive pediatric brain tumor growth.

---

**Extended Data Fig. 1: ZFTA-RELA fusion regulates metabolic programs associated with methionine metabolism.** (A) Schematic illustrating the establishment of patient-derived supratentorial ependymoma (ST-EPN) cell lines from primary tumors.

(B) Representative immunoblots showing expression of the ZFTA-RELA fusion protein and endogenous RELA across patient-derived ST-EPN cell lines. A ZFTA-MAML3 fusion-positive ST-EPN cell line was included as a control. ZR^Fus^, ZFTA-RELA fusion-positive. Vinculin served as the loading control.

(C) Representative immunoblots showing expression of the ZFTA-RELA fusion protein in two independent Cas9-expressing patient-derived ZR^Fus^ cell lines following transduction with control single-guide RNA (sgRNA; sgCONT) or sgRNA targeting *ZFTA-RELA* (sgZFTA-RELA). Vinculin served as the loading control.

(D) Cell proliferation of ZR^Fus^ cell lines following transduction with sgCONT or sgZFTA-RELA. Cell number was measured over 5 days.

(E) Integrated metabolomic and transcriptomic pathway analysis following ZFTA-RELA knockout (KO) in ZR^Fus^ cells, incorporating significantly depleted metabolites (p ≤ 0.05 and log₂ fold change ≤ −0.58) and downregulated genes (false discovery rate (FDR) ≤ 0.05 and log₂ fold change ≤ −1). Methionine metabolism was among the suppressed pathways.

(F) Dot plot showing metabolic pathway-associated gene expression across annotated cell populations from single-cell RNA sequencing (scRNA-seq) of primary patient ZR^Fus^ ST-EPN tumors. Dot size represents the percentage of cells expressing pathway-associated genes, and color indicates average expression across each cell population.

For panel D, data are presented as mean ± SD from three independent samples per condition. For panel E, metabolomic and transcriptomic analyses were performed using three independent samples per group. Panel F comprises scRNA-seq data from four primary ZR^Fus^ ST-EPN tumors. Statistical significance was determined using two-way ANOVA with Šidák’s multiple-comparisons test for panel D. ****p < 0.0001.

**Extended Data Fig. 2: ZFTA-RELA fusion regulates transcriptional programs associated with methionine metabolism.** (A) Heatmaps showing differentially expressed genes (DEGs; |log₂ fold change| ≥ 1 and FDR ≤ 0.05) following ZFTA-RELA knockout (KO) in two independent patient-derived ZR^Fus^ supratentorial ependymoma (ST-EPN) cell lines.

(B) Venn diagram showing overlap of DEGs following ZFTA-RELA KO in ZR^Fus^ cell lines. The accompanying bar plot shows the number of commonly upregulated and downregulated genes.

(C) Top downregulated pathways identified by Gene Set Enrichment Analysis (GSEA) following ZFTA-RELA KO. RNA sequencing (RNA-seq) data from two independent ZR^Fus^ cell lines were combined for pathway analysis. Pathways with FDR q ≤ 0.25 were considered significant.

(D) Schematic illustrating ectopic expression of ZFTA-RELA in embryonic day 13.5 (E13.5) mouse neural stem cells (mNSCs).

(E) Representative immunoblots showing expression of HA-tagged ZFTA-RELA fusion protein and endogenous RELA in mNSCs expressing empty vector (EV) or ZFTA-RELA. Vinculin served as the loading control.

(F) Cell proliferation of EV- and ZFTA-RELA-expressing mNSCs. Cell number was measured over 5 days.

(G) Heatmap showing DEGs (|log₂ fold change| ≥ 1 and FDR ≤ 0.05) following ZFTA-RELA overexpression (OE) in mNSCs. The accompanying bar plot shows the numbers of upregulated and downregulated genes.

(H) Top enriched pathways identified by GSEA following ZFTA-RELA OE in mNSCs. Pathways with FDR q ≤ 0.25 were considered significant.

(I) Cross-species comparison of pathways downregulated following ZFTA-RELA KO in patient-derived ZR^Fus^ cells and upregulated following ZFTA-RELA OE in mNSCs.

(J) Reciprocal pathway enrichment analysis comparing ZFTA-RELA KO and OE RNA-seq datasets. Dot size represents −log₁₀(FDR), and the x-axis indicates the normalized enrichment score (NES).

(K) GSEA enrichment plots showing significant downregulation of methionine metabolism following ZFTA-RELA KO (NES = −1.51, FDR q = 0.033) and enrichment following ZFTA-RELA OE (NES = 1.61, FDR q = 0.018).

For panel F, data are presented as mean ± SD from three independent samples per condition. For panels A–C and G–H, RNA-seq was performed using three independent samples per group. Statistical significance was determined using two-way ANOVA with Šidák’s multiple-comparisons test for panel F. ***p < 0.001, ****p < 0.0001.

**Extended Data Fig. 3: AHCY regulates tumor growth, methionine-cycle activity, and pyrimidine metabolism in ZFTA-RELA fusion ependymoma.** (A) Uniform Manifold Approximation and Projection (UMAP) visualization of scRNA-seq data from in utero electroporation (IUE)-derived murine ZR^Fus^ tumors (12,179 cells). Each dot represents an individual cell and is color-coded according to cell identity. Feature plots show expression of *Slc7a5* and *Slc43a2* across individual cell populations.

(B) Chromatin immunoprecipitation sequencing (ChIP-seq) occupancy of HA-tagged ZFTA-RELA at the *Slc7a5* locus in murine ZR^Fus^ supratentorial ependymoma (ST-EPN) tumors. Immunoglobulin G (IgG) served as the negative control. Data were obtained from Arabzade *et al.* (*Cancer Discovery*, 2021; GEO: GSE161679).

(C) Relative abundance of S-adenosylmethionine (SAM) in patient-derived ZFTA-MAML3 fusion-positive and ZR^Fus^ ST-EPN cells.

(D) Cell proliferation of three independent patient-derived ZR^Fus^ cell lines expressing control shRNA (shCONT) or shRNA targeting AHCY (shAHCY). Cell number was measured over 5 days.

(E) Cell number of two patient-derived ZR^Fus^ cell lines following treatment with vehicle, the AHCY inhibitor aristeromycin (2.5 μM), or aristeromycin supplemented with homocysteine (100 μM).

(F) Heatmaps showing differentially expressed genes (DEGs; |log₂ fold change| ≥ 1 and false discovery rate (FDR) ≤ 0.05) following AHCY knockdown (KD) in two independent patient-derived ZR^Fus^ cell lines.

(G) Venn diagram showing overlap of DEGs following AHCY KD in ZR^Fus^ cell lines. The accompanying bar plot shows the numbers of commonly upregulated and downregulated genes.

(H) Metabolic pathway enrichment analysis of significantly depleted metabolites (n = 40) following AHCY KD in patient-derived ZR^Fus^ cells.

(I) Relative abundance of N-carbamoyl-L-aspartate and dihydroorotate in murine ZR^Fus^ tumors compared with normal cortex.

(J) Relative abundance of N-carbamoyl-L-aspartate and dihydroorotate in patient-derived ZR^Fus^ cells following CRISPR/Cas9-mediated ZFTA-RELA knockout compared with control single-guide RNA (sgRNA; sgCONT).

(K) Representative immunoblots showing ASNS expression in pediatric high-grade glioma (Ped-HGG) cell lines following AHCY KD. Vinculin served as the loading control.

For panels C-E and J, data are presented as mean ± SD from three independent samples per group or condition. For panel I, data are presented as mean ± SD from four biological replicates per group. Panel A comprises scRNA-seq data from two murine IUE-derived ZR^Fus^ tumors. For panels F and G, RNA-seq was performed using three independent samples per group. For panel H, metabolomic analysis was performed using three independent samples per group. Statistical significance was determined using an unpaired two-tailed t-test with Welch’s correction for panels C, I, and J, two-way ANOVA with Šidák’s multiple-comparisons test for panel D, and one-way ANOVA with Šidák’s multiple-comparisons test for panel E. *p < 0.05, **p < 0.01, ****p < 0.0001; ns, not significant.

**Extended Data Fig. 4: AHCY regulates ASNS expression independently of the integrated stress response in ZFTA-RELA fusion ependymoma.** (A) H3K4me3 chromatin immunoprecipitation quantitative PCR (ChIP-qPCR) analysis at the CAD promoter/transcription start site (TSS) in patient-derived ZFTA-RELA fusion-positive (ZR^Fus^) supratentorial ependymoma (ST-EPN) cells expressing control shRNA (shCONT) or shRNA targeting AHCY (shAHCY). Immunoglobulin G (IgG) served as the negative control.

(B) Schematic illustrating regulation of ASNS through the integrated stress response (ISR). Amino acid depletion or asparagine limitation activates GCN2 and eIF2α phosphorylation, leading to ATF4-dependent transcriptional activation of ASNS. GCN2, general control nonderepressible 2; eIF2α, eukaryotic translation initiation factor 2 alpha; ATF4, activating transcription factor 4.

(C) Representative immunoblots showing phosphorylated GCN2 (p-GCN2; Thr899), phosphorylated eIF2α (p-eIF2α; Ser51), and ATF4 levels in two independent patient-derived ZR^Fus^ cell lines expressing shCONT or shAHCY. Vinculin served as the loading control.

(D) Densitometric quantification of ATF4 protein expression following AHCY knockdown in ZR^Fus^ cell lines, normalized to shCONT.

(E) Representative immunoblots showing ASNS expression in two independent patient-derived ZR^Fus^ ST-EPN cell lines following treatment with dimethyl sulfoxide (DMSO) or the G9a/GLP H3K9 methyltransferase inhibitor UNC0638 for 72 h. Vinculin served as the loading control.

(F) Representative immunoblots showing ASNS expression in Jurkat, MOLT-4, and RS4;11 cells following treatment with dimethyl sulfoxide (DMSO) or the EZH1/2 inhibitor UNC-1999 (2 μM, 72 h). Vinculin served as the loading control.

(G) Representative immunohistochemistry (IHC) images showing ASNS expression in human diffuse intrinsic pontine glioma patient-derived xenograft (DIPG PDX) and medulloblastoma patient-derived xenograft (MB PDX) tissues. Scale bar, 50 μm.

(H) Normalized cell number at day 5 in KNS42 and DIPG-07 cells following ASNS overexpression (ASNS OE) compared with empty vector (EV) control.

(I) Volcano plot showing differential metabolites in patient-derived ZR^Fus^ cells following ASNS overexpression (ASNS OE) compared with empty vector (EV). Significantly altered metabolites are highlighted in red (enriched) and blue (depleted). Aspartate, N-carbamoyl-L-aspartate, and asparagine are indicated.

For panels A, D and H, data are presented as mean ± SD from three independent samples per condition. For panel I, metabolomic analysis was performed using three independent samples per group. Statistical significance was determined using an unpaired two-tailed t-test with Welch’s correction for panels A, D, and H. *p < 0.05; ns, not significant.

**Extended Data Fig. 5: ZFTA-RELA fusion ependymoma upregulates SLC1A5 expression and is sensitive to asparagine depletion.** (A) Relative cell number of two in utero electroporation (IUE)-derived murine ZFTA-RELA fusion-positive (ZR^Fus^) ependymoma cell lines (left) and two diffuse intrinsic pontine glioma (DIPG) cell lines (right) cultured with 10 μM or 100 μM asparagine. Values were normalized to the corresponding 10 μM asparagine condition.

(B) Dot plot showing *SLC1A5* and *ASNS* expression across cell populations identified by single-cell RNA sequencing (scRNA-seq) of primary human ZR^Fus^ tumors. Dot size represents the percentage of cells expressing each gene, and color indicates average expression.

(C) Dot plot showing *Slc1a5* and *Asns* expression across cell populations identified by scRNA-seq of IUE-derived murine ZR^Fus^ tumors. Dot size represents the percentage of cells expressing each gene, and color indicates average expression.

(D) ZFTA-RELA-HA chromatin immunoprecipitation sequencing (ChIP-seq) occupancy at the *Slc1a5* locus in murine ZR^Fus^ supratentorial ependymoma (ST-EPN) tumors. Immunoglobulin G (IgG) served as the negative control. Data were obtained from Arabzade *et al.* (*Cancer Discovery*, 2021; GEO: GSE161679).

(E) H3K27ac ChIP-seq tracks at the *SLC1A5* locus in three patient-derived ST-EPN tumors. Data were obtained from Mack *et al.* (2017).

(F) Representative brightfield images of patient-derived ZR^Fus^ and ZFTA-MAML3 fusion-positive ST-EPN cells following treatment with vehicle or L-asparaginase (ASNase; 1 U ml⁻¹). Scale bar, 100 μm.

(G) Representative immunoblots showing ASNS expression in two independent patient-derived ZR^Fus^ cell lines following treatment with vehicle or ASNase (1 U ml⁻¹). Vinculin served as the loading control. ASNS levels relative to vinculin are indicated.

(H) Dose-response curves showing viability of patient-derived (left) and IUE-derived murine (right) ZR^Fus^ cell lines following 96 h treatment with increasing concentrations of the ASNS inhibitor ASX-173 in the presence of vehicle or ASNase (1 U ml⁻¹).

(I) Representative brightfield images of patient-derived ZR^Fus^ cells and ZFTA-RELA-expressing mouse neural stem cells (mNSCs) following treatment with vehicle, ASNase (1 U ml⁻¹), or ASNase supplemented with aspartate (10 mM). Scale bar, 100 μm.

For panels A, and H, data are presented as mean ± SD from three independent samples per condition. Statistical significance was determined using an unpaired two-tailed t-test with Welch’s correction for panel A. ***p < 0.001, ****p < 0.0001; ns, not significant.

**Extended Data Fig. 6: ZFTA-RELA fusion ependymoma is sensitive to DHODH inhibition.** (A) Comparison of isotopologue fractions of aspartate and N-carbamoyl-L-aspartate in patient-derived ZFTA-RELA fusion-positive (ZR^Fus^) supratentorial ependymoma (ST-EPN) cells.

(B) Schematic illustrating inhibition of de novo pyrimidine biosynthesis by BAY-2402234 through targeting dihydroorotate dehydrogenase (DHODH).

(C) Dose-response curves showing viability of three patient-derived ZR^Fus^ ST-EPN cell lines (left), three glioma cell lines (middle), and three diffuse intrinsic pontine glioma (DIPG) cell lines (right) following 72 h treatment with increasing concentrations of BAY-2402234.

(D) Representative brightfield images of patient-derived ZR^Fus^ cells (top) and ZFTA-RELA-expressing mouse neural stem cells (mNSCs; bottom) following treatment with vehicle, BAY-2402234 (2.5 nM), or BAY-2402234 plus uridine (0.1 mM). Scale bar, 100 μm.

(E) Representative immunoblots showing glycosylated SLC1A5 expression in two independent patient-derived ZR^Fus^ cell lines following ASNS overexpression (ASNS OE) compared with EV control. Vinculin served as the loading control.

(F) Representative hematoxylin and eosin (H&E)-stained coronal brain sections from mice bearing orthotopic tumors generated from ZFTA-RELA-expressing mNSCs (top) or in utero electroporation (IUE)-derived ZR^Fus^ tumors (bottom) following treatment with vehicle or BAY-2402234.

For panels A and C, data are presented as mean ± SD from three independent samples per condition. Statistical significance was determined using an unpaired two-tailed t-test with Welch’s correction for panel A. ns, not significant.

## Acknowledgements

K.A.M. is supported by the Matthew Larson Foundation (IronMatt) AWD00012909, the Stanley M. Marks Cancer Research Fund, the V Foundation for Cancer Research GC2025-007, UPMC-Children’s Hospital Nyssen Fund, Walter L. Copeland Fund by the Pittsburgh foundation, Children’s Neuroscience Institute-Interdisciplinary Neuroscience Award (INA), Children’s Trust Grantmakers’ Award, and UPMC Hillman Cancer Center core grant (NCI, P30CA047904). N.Y. is supported by the Walter L. Copeland Fund by the Pittsburgh foundation. R.B.C. is supported by T32. S.V. is supported by R01NS110572, R01NS127799, and R01CA261926. We thank the Animal Imaging Core Laboratory at UPMC Children’s Hospital of Pittsburgh for assistance with animal imaging studies. Metabolomics services were performed by the Metabolomics Core Facility at Robert H. Lurie Comprehensive Cancer Center (generously supported by NCI CCSG P30 CA060553) of Northwestern University. This project used the UPMC Hillman Cancer Center and Tissue and Research Pathology/Pitt Biospecimen Core shared resource which is supported in part by award P30CA047904.

## Author contributions

N.Y. and K.A.M. conceived and designed the study, analyzed and interpreted the data, generated the figures, and wrote the manuscript. O.P., R.G.R.M., Senna Visser, A.C.T.C., and B.M. contributed to immunohistochemistry experiments. R.G.R.M. performed ChIP experiments and associated data analysis. K.B.N.R., B.M., V.P., Y.K., and S.D.V. contributed to immunoblotting experiments. J.Z., A.Y.M., and S.R.K. contributed to single-cell RNA-sequencing data analysis. S.K.N. performed ATAC-seq experiments and associated data analysis. Y.P. contributed to animal experiments.

R.B.C. and H.S. contributed to metabolomic experiments and data analysis. A.D., K.G.A., B.H., C.-H.G.L., P.S., F.S., A.M.D., N.K.F., M.F.R., T.M.P., P.D., K.W.P., S.C.M., Sriram Venneti, J.R., R.J.G., E.C.H., S.K.R., T.J.A., S.R., S.A., B.Z., B.F., M.D.T., E.H., I.F.P., and T.P. provided resources for the study. K.A.M. supervised the study and secured funding. All authors reviewed and approved the manuscript.

## Experimental models

### Primary patient tumor tissue

Primary patient-derived supratentorial ependymoma (ST-EPN) tissues analyzed in this study were obtained as part of a previously published study^21^. Tumor specimens were collected at the time of surgical resection from patients who had not received radiotherapy or chemotherapy before surgery.

### Mouse models

All animal experiments were conducted in accordance with protocols approved by the Institutional Animal Care and Use Committee (IACUC) of the University of Pittsburgh (IACUC Protocol 25066598). Male and female C57BL/6J mice (The Jackson Laboratory, stock no. 000664), 4–6 weeks of age, were used for intracranial implantation of in utero electroporation (IUE)-derived murine ependymoma cells. NOD-SCID-IL2Rγ-chain-deficient (NSG) mice (NOD.Cg-*Prkdc*scid*Il2rg*tm1Wjl/SzJ); The Jackson Laboratory, stock no. 005557), 4–6 weeks of age, were used for orthotopic implantation of ZFTA–RELA-expressing murine neural stem cells (mNSC_ZR^Fus^). J:NU nude mice (The Jackson Laboratory, stock no. 007850), 4 weeks of age, were used for subcutaneous flank tumor implantation experiments. Mice were maintained under a 14-h light/10-h dark cycle at an ambient temperature of 18–23 °C, with no more than five mice housed per cage. Animal health and housing conditions were routinely monitored by animal husbandry staff under veterinary supervision.

### In utero electroporation-derived tumors

Murine ZFTA–RELA fusion-positive ependymomas (mZR^Fus^-EPN) were generated by in utero electroporation (IUE) of embryonic day 13.5–14.5 C57BL/6J mouse embryos, as previously described^26^. Briefly, approximately 1 µL of a plasmid and mRNA mixture containing 0.05% Fast Green was injected into the lateral ventricle of each embryo. The mixture included the piggyBac-based constructs PBCAG-ZFTA–RELA-HA, PBCAG-eGFP, and pCAG-PBase, together with Cas9 mRNA and sgRNAs targeting *Cdkn2a*. Electroporation was performed using five 45-V square pulses of 50 ms duration separated by 950-ms intervals, with 3-mm tweezer electrodes directed toward the dorsal cortex. Successfully electroporated offspring were monitored for tumor development and euthanized upon the onset of neurological symptoms associated with tumor burden. All procedures were performed at the University of Cincinnati under approved IACUC protocols 22-05-15-01 and 25-05-05-01.

### Cell lines

#### Human cell lines

Primary patient-derived ST-EPN cell lines harboring the ZFTA–RELA fusion included ST-02, ST-03, ST-04, CP-01 (CPITT-1), and EP-01, whereas ST-01 harboured the ZFTA–MAML3 fusion. ST-02, ST-04, and ST-01 were established and characterized as previously described^21^. EP1NS (EP-01) was provided by S.C.M., and CPITT-01 (CP-01) was provided by S.A.

Human fetal neural stem cells (hfNSCs; HF7450) were obtained from P.D. Posterior fossa group A (PFA) ependymoma cell lines included PFA-09, EP-06, and EPD-210 were maintained as previously described^21^. Pediatric glioma cell lines KNS42, SF-188, and SJ-G2 were provided by Dr. I.F.P. Diffuse intrinsic pontine glioma (DIPG) cell lines DIPG-04, DIPG-06, DIPG-07, and DIPG-13 were provided by Dr. Michelle Monje (Stanford University, Stanford, CA, USA). Acute lymphoblastic leukemia (ALL) cell lines Jurkat, MOLT-4, and RS4;11 was provided by Dr. Wei Du (UPMC Hillman Cancer Center, Pittsburgh, PA, USA). The D425 medulloblastoma cell line was provided by Dr. M.D.T. (Texas Children’s Hospital, Houston, TX, USA).

#### Mouse cell lines

Control murine neural stem cells (mNSCs) were derived from the forebrain of embryonic day 13 (E13) C57BL/6J mouse embryos, as previously described^21^. The ZFTA–RELA-expressing murine neural stem cell line (mNSC_ZR^Fus^) was provided by S.V. In utero electroporation (IUE)-derived murine ependymoma cell lines ZR^Fus^Line_IUE-01 and ZR^Fus^Line_IUE-05 were provided by T.P.

### Methods

#### Cell culture conditions

Primary patient-derived ST-EPN cell lines, hfNSCs, and PFA ependymoma cell lines were cultured on plates sequentially coated with poly-L-ornithine (Sigma #P4957) and laminin (Sigma #L2020) as previously described^21,68^. Cells were maintained in serum-free Neurobasal medium (Thermo Fisher Scientific #3137184) supplemented with N-2 supplement (Thermo Fisher Scientific #17502-048), B-27 supplement minus vitamin A (Thermo Fisher Scientific #12587-010), EGF (10 ng mL⁻¹; PeproTech #AF-100-15), FGF2 (10 ng mL⁻¹; PeproTech #100-18C), heparin (2 µg mL⁻¹; Sigma #H3393), bovine serum albumin (0.015%; Sigma #A8412), GlutaMAX (1×; Thermo Fisher Scientific #35050061), and Antibiotic–Antimycotic (1×; Thermo Fisher Scientific #15240062). Culture medium was refreshed every 2–3 days. Cells were passaged at approximately 70–80% confluence using Accutase (Thermo Fisher Scientific #A1110501), diluted with DMEM (Thermo Fisher Scientific #11995065) at a 1:3–1:4 ratio, and replated onto freshly coated plates. Pediatric glioma cell lines were maintained in DMEM (Thermo Fisher Scientific #11995065) supplemented with 10% heat inactivated fetal bovine serum (FBS; Thermo Fisher Scientific #16140071) and Antibiotic–Antimycotic (1×). DIPG cells were maintained in DIPG culture medium as previously described^19^. ALL cell lines were maintained as previously described^69^. Mouse cell lines were maintained in serum-free complete NeuroCult™ Proliferation Medium (STEMCELL Technologies #05702) supplemented with EGF (10 ng mL⁻¹; PeproTech #315-09), FGF2 (10 ng mL⁻¹; PeproTech #450-33), heparin (2 µg mL⁻¹), and Antibiotic–Antimycotic (1×).

All cell cultures were maintained at 37 °C in a humidified incubator with 5% CO₂. Cell line identities were confirmed by short tandem repeat (STR) profiling, and cultures were routinely tested for mycoplasma contamination by PCR upon thawing of new cell stocks and at least annually thereafter. Early- to mid-passage cells, fewer than 20 passages, were used for experiments.

### Metabolite extraction and mass spectrometry

#### Human tissue

Global metabolite abundance profiling of snap-frozen ST-EPN tissues and age-matched normal cortex tissues was performed in collaboration with Metabolon, Inc., as part of a previously published study^21^. Metabolite extraction, sample preparation, mass spectrometry, metabolite identification, and data processing were performed according to Metabolon’s established protocols.

#### Mouse tissue

Mouse tumor tissues and adjacent normal cortex tissues were pulverized under liquid nitrogen using a pre-chilled mortar and pestle to prevent sample thawing. Pulverized tissues were accurately weighed into pre-chilled screw-cap tubes (Sarstedt #NC9180825) and stored at −80 °C until metabolite extraction and liquid chromatography–mass spectrometry (LC–MS) analysis. Tissue samples were extracted with acetonitrile:Methanol:water, 2:2:1 v/v/v containing 0.1 M formic acid at fixed ratio of 20μL per mg tissue, quickly vortexed and after 3 mins, it was neutralized with 15% ammonium bicarbonate (8.7 µL per 100 µL solvent). The homogenate was processed using a bead mill homogenizer (Omni International), sonicated at 0 °C and subjected to a freeze–thaw cycle; these steps were repeated once. Following incubation at 0 □°C for 20 □min, samples were centrifuged at 18,000g for 20 □min at 4 □°C. The supernatant was collected, and aliquots were immediately subjected to LC–MS analysis. A pooled quality control (QC) sample was prepared by combining equal volumes of supernatant from each sample. For LC–MS analysis, metabolite separation and detection were performed using a Thermo Scientific Vanquish Horizon UHPLC system coupled to an Orbitrap IQ-X Tribrid mass spectrometer (Thermo Scientific). Chromatographic separation was achieved on an iHILIC-(P) Classic column (2.1 × 150 □mm, 5 □µm; part no. 160.152.0520; HILICON). The mobile phase consisted of 20 mM ammonium bicarbonate (pH 9.6, adjusted with ammonium hydroxide) as mobile phase A (MPA) and acetonitrile as mobile phase B (MPB). The column temperature was maintained at 40 □°C, with an injection volume of 2 □µl and a flow rate of 0.2 ml min−1. The chromatographic gradient was: 0 □min, 85% B; 0.5 □min, 85% B; 18 □min, 20% B; 20 min, 20% B; 20.5 min, 85% B; and 28 min, 85% B. MS was performed in both positive- and negative ionization modes, with spray voltages of 3,600 V and 2,800 V, respectively. Other parameters were: sheath gas, 35 arb; auxiliary gas, 5 arb; sweep gas, 1 arb; ion-transfer-tube temperature, 250 □°C; vaporizer temperature, 350 □°C; AGC target, 100%; and maximum injection time, 118 □ms. Data were acquired in full-scan mode (m/z 70–1,000) at ×60,000 resolution using the Xcalibur software package (Thermo Scientific). Metabolite identification was performed by matching accurate mass (±5 ppm), retention time, and, when available, MS/MS fragmentation spectra to reference standards in an in-house library. Data processing was performed using the Compound Discoverer 3.3(Thermo Scientific) or Skyline^70^.

#### Cell lines

Hydrophilic metabolites were extracted from cultured cells as previously described^71^. On the day of extraction, an additional plate of cells was harvested in parallel to determine cell numbers for sample normalization. Following aspiration of the culture medium, cells were washed twice with ice-cold saline to remove residual medium and immediately placed on dry ice. Cellular metabolism was quenched by adding 1 mL of pre-chilled 80% methanol (v/v; −80 °C) directly to each plate. Cells were scraped, and the resulting lysates were transferred to pre-chilled microcentrifuge tubes. Samples were subjected to three freeze–thaw cycles between −80 °C and room temperature, with brief vortexing between cycles, to ensure complete cell lysis. Lysates were subsequently incubated overnight at −20 °C to precipitate proteins and centrifuged at 20,000 × g for 15 min at 4 °C. The clarified supernatants containing polar metabolites were collected, and extracts normalized to equivalent cell numbers were submitted to the Northwestern University Metabolomics Core Facility for analysis. Dried metabolite extracts were reconstituted in 60% acetonitrile (v/v), vortexed for 30 s, and centrifuged at 20,000 × g for 30 min at 4 °C to remove insoluble material. The resulting supernatants were subjected to LC–MS analysis. Metabolite profiling was performed using high-performance liquid chromatography coupled to high-resolution tandem mass spectrometry (HPLC–MS/MS) on a Thermo Q Exactive mass spectrometer equipped with an electrospray ionization (ESI) source and coupled to an Ultimate 3000 HPLC system (Thermo Fisher Scientific). Metabolites were separated using a Waters XBridge Amide column (3.0 mm × 100 mm, 3.5 µm particle size). Mobile phase A consisted of 95% water, 5% acetonitrile, 10 mM ammonium hydroxide, and 10 mM ammonium acetate (pH 9.0), whereas mobile phase B consisted of 100% acetonitrile. The gradient elution program was as follows: 0 min, 15% A; 2.5 min, 64% A; 12.4 min, 40% A; 12.5– 14 min, 30% A; and 14–21 min, 15% A, at a flow rate of 150 µL min⁻¹. The ESI source was operated at 4.0 kV with a capillary temperature of 275 °C, sheath gas setting of 35 arbitrary units, and auxiliary gas setting of 5 arbitrary units. Data were acquired in polarity-switching mode over an m/z range of 60–900. MS¹ spectra were acquired at a resolution of 70,000 with an automatic gain control (AGC) target of 1 × 10 □and a maximum injection time of 200 ms. MS² spectra were acquired using higher-energy collisional dissociation (HCD) at a normalized collision energy of 30% and a resolution of 17,500. Metabolites were identified based on accurate mass-to-charge ratio (m/z), retention time matching to authentic standards, and/or MS² fragmentation patterns. Data acquisition and processing were performed using Xcalibur 4.1 and TraceFinder 4.1 software (Thermo Fisher Scientific).

### U-¹³C_₆_-glucose and amide-¹□N-glutamine isotope tracing

Tracing medium was prepared using serum-free DMEM lacking glucose, glutamine, and phenol red (Thermo Fisher Scientific #A1443001) and supplemented with N-2 supplement, B-27 supplement minus vitamin A, EGF, FGF2, heparin, bovine serum albumin, and Antibiotic–Antimycotic (1×) at the concentrations described above. Stable isotope tracing experiments were performed for 8 h in separate plates using either U-¹³C₆-glucose or amide-¹ N-glutamine. On the day of labeling, cells maintained in complete serum-free DMEM were washed twice with calcium- and magnesium-free phosphate-buffered saline (PBS) to remove residual unlabeled nutrients. For glucose tracing, cells were incubated for 8 h in tracing medium containing 25 mM U-¹³C₆-glucose (Cambridge Isotope Laboratories #CLM-1396) and 2 mM unlabeled glutamine. For glutamine tracing, cells were incubated for 8 h in tracing medium containing 25 mM unlabeled glucose and 2 mM amide-¹ N-glutamine (Cambridge Isotope Laboratories #NLM-557). Following labeling, the medium was aspirated, and cells were rapidly washed twice with ice-cold calcium- and magnesium-free PBS to remove residual extracellular tracer. Plates were immediately placed on dry ice, and metabolites were extracted and processed as described above for steady-state metabolite analysis. Metabolite extracts normalized to equivalent cell numbers were submitted to the Northwestern University Metabolomics Core Facility for LC–MS analysis.

### U-¹³C_4_-asparagine uptake

For asparagine uptake experiments, cells maintained in complete culture medium were washed to remove residual extracellular asparagine and incubated for 8 h with 0.1 mM U-¹³C₄-asparagine (Cambridge Isotope Laboratories #CLM-8699-H) in the presence of either DMSO or the SLC1A5 inhibitor V-9302. Following labeling, 500 μl of culture supernatant was collected and centrifuged at 600g for 5 min at room temperature to remove cellular debris. A 300-μl aliquot of cleared supernatant was collected and stored at −80 °C until LC–MS analysis. For intracellular metabolite extraction, the remaining medium was aspirated and cells were gently washed with 0.9% (w/v) saline at room temperature to remove residual extracellular tracer. Plates were immediately transferred to ice to quench metabolism, and metabolites were extracted using dry ice-cold 80% methanol-based extraction solvent. Samples were incubated at −80 °C for 30 min, cells were scraped, and extracts were transferred to screw-cap tubes and shipped on dry ice to University of Chicago Comprehensive Cancer Metabolomics Platform. The extract was thaw on ice, centrifuged at 18,000g and 4°C for 20 mins, supernatant was dried and re-suspended in 60/40 acetonitrile/water before the analysis. The LC-MS method was the same as the mouse tissue profiling as described earlier for the intracellular labeled asparagine detection. For the media asparagine detection, 30 μl of media was mixed with the 120 μl of acetonitrile, vortex, incubate on ice, centrifuge at 18,000g and 4°C and the supernatant was injected on BEH Z-HILIC (2.1×150 mm, 2.5 µM; part # 186009990; Waters Corporation) column at acidic pH. The mobile phase A (MPA) was 10 mM ammonium formate containing 0.2% formic acid and mobile phase B (MPB) was acetonitrile containing 0.1% formic acid. The column temperature, injection volume, and flow rate were 30°C, 5 µL, and 0.2 mL/minute, respectively. The chromatographic gradient was 0 minute: 90% B, 15 minutes: 20% B, 16 minutes: 20% B, 16.5 minutes: 90% B, 17 minutes: 90% B, and 23 minutes: 90% B. During the re-equilibration, the flow rate was increased to 0.4mL/minute for 4.7 minutes. MS detection was done using Orbitrap IQ-X Tribrid mass spectrometer (Thermo Scientific) with a H-ESI probe operating in switch polarity mode for both methods. MS parameters were as follows: spray voltage: 3800 V for positive ionization and 2500 V for negative ionization modes, sheath gas: 80, auxiliary gas: 25, sweep gas: 1, ion transfer tube temperature: 300°C, vaporizer temperature: 300°C, automatic gain control (AGC) target: 25%, and a maximum injection time of 80 milliseconds (ms). The data acquisition was done using the Xcalibur software (Thermo Scientific) in full-scan mode with a range of 70-600 m/z at 120K resolution. Data processing was performed using the Skyline software and natural abundance correction was performed using the IsoCorrectoR^72^.

### Metabolomic data analysis

Metabolomic data were analyzed using MetaboAnalyst 6.0. Metabolite enrichment and pathway analyses were performed using annotated metabolites to identify significantly altered metabolic processes and pathways between experimental groups. For integrative analysis, joint pathway analysis was performed by combining metabolomic and RNA-sequencing (RNA-seq) data to identify pathways coordinately altered at the metabolite and gene-expression levels. Pathway enrichment and topology analyses were used to prioritize biologically relevant metabolic pathways. Volcano plots of differential metabolite abundance were generated in R using the ggplot2 package.

### Single-cell RNA sequencing and analysis of primary patient and IUE-derived tumors

Primary patient ST-EPN tissues were obtained as part of a previously published study^21^. Briefly, freshly resected tumors were mechanically and collagenase-dissociated to generate single-cell suspensions. Approximately 10,000–15,000 cells per sample were loaded onto the 10x Genomics Chromium platform using Single Cell 3′ v3/v3.1 chemistry, and libraries were sequenced on an Illumina NovaSeq 6000. ST-EPN scRNA-seq reads were aligned to the human GRCh38 reference (v2024-A; 10x Genomics) using Cell Ranger v10.0.0 with default settings to generate gene–barcode count matrices. ST-EPN scRNA-seq samples were aligned to the human genome reference GRCh38 (v2024-A; 10x Genomics) using the count pipeline in CellRanger (v10.0.0) with default analysis settings to generate raw gene–barcode count matrices. Quality control was performed separately for each sample using R package Seurat (v5.4.0) before normalization and integration. Genes detected in at least 10 cells and cells with at least 500 detected genes were retained. Sample-specific filtering thresholds were calculated using the first quartile (Q1), third quartile (Q3), and interquartile range (IQR) of each quality control metric. Cells were retained if both the number of detected genes and total UMI counts were greater than Q1 − 1.5 × IQR and less than Q3 + 3 × IQR, and if the percentage of mitochondrial counts was below Q3 + 3 × IQR. Doublets were identified and removed using R package DoubletFinder (v2.0.3), with the expected doublet rate based on the multiple rate provided by 10x Genomics. Individual samples were then merged and jointly normalized using R package SCTransform (v0.4.3), with variable features, n = 3000, while regressing out variation associated with mitochondrial count percentage and cell-cycle scores. Clustering analysis was performed using Seurat (v5.4.0). Principal component analysis (PCA) was used for linear dimensionality reduction to identify statistically significant principal components. Harmony (v1.2.4), with theta = 2, was applied to account for sample-associated variation. Cell clusters were identified using Seurat’s shared nearest neighbor (SNN) modularity optimization algorithm and visualized using uniform manifold approximation and projection (UMAP). Cell types were manually annotated based on established cell-type marker genes and differentially expressed genes (DEGs) identified using Seurat’s FindAllMarkers function with default parameters. Marker genes for each cluster are provided in Supplementary Table 12.

For IUE-derived ST-EPN tumors, approximately 100,000 sorted cells per sample were fixed using the Evercode Low Input Fixation Kit v4 (Parse Biosciences #LCF500) following the manufacturer’s protocol. Fixed cells were subsequently barcoded and prepared for sequencing with the Evercode Mega v4 kit (ECWT4500). Final libraries were sequenced on an Illumina NovaSeq X Plus platform using paired-end reads, targeting approximately 20,000 reads per cell. Sequencing data were demultiplexed, aligned to the mm10 reference genome, and processed to generate gene-count matrices using the Parse Biosciences Trailmaker pipeline.

### Lentiviral packaging and infection

To generate ZFTA-RELA fusion knock-out cell lines, single-guide RNA (sgRNA) targeting the ZFTA exon 3 sequence at the ZFTA–RELA fusion breakpoint (5′-CTCATGGACTATGACCCGCG-3′) or a non-targeting control sgRNA (5′-GCACTACCAGAGCTAACTCA-3′) were designed and purchased from VectorBuilder. sgRNAs were delivered together with a Cas9 expression vector (lentiCRISPR v2; Addgene #52961) into cells via lentiviral transduction, as described below. Cells were selected with puromycin and expanded as pooled populations. Fusion disruption was confirmed by immunoblotting for ZFTA-RELA protein. To generate isogenic murine neural stem cell (mNSC) models, cells were transduced with lentiviral particles encoding ZFTA-RELA or empty vector (EV) control. Stable expression was established by antibiotic selection and confirmed by immunoblotting.

For stable knockdown experiments, shRNA constructs targeting human *AHCY* (TRCN0000229329 and TRCN0000229330), mouse *Ahcy* (TRCN0000302750 and TRCN0000349016), mouse *Ezh2* (TRCN0000301834 and TRCN0000304506), mouse *Slc7a5* (TRCN0000079395 and TRCN0000079396), and mouse *Slc1a5* (TRCN0000079258 and TRCN0000327527) were obtained from Sigma-Aldrich. For overexpression studies, human *ASNS* constructs and empty vector (EV) controls were purchased from VectorBuilder; *ZFTA–RELA* fusion construct was provided by S.C.M and K.W.P.; and *EZHIP* construct was provided by Dr. Peter W Lewis, University of Wisconsin-Madison, Madison, WI, US. A scrambled shRNA control (Addgene plasmid #1864) was used as a non-targeting control. All constructs were delivered via lentiviral transduction, and stable cell populations were generated by antibiotic selection for downstream assays.

For lentivirus production, shRNA and overexpression plasmids were propagated in *Escherichia coli* cultured in LB medium and purified using the ZymoPURE™ II Plasmid Midiprep Kit (Zymo Research #NC0835048). Plasmid DNA concentrations were determined by absorbance at 260 nm using a NanoDrop One spectrophotometer. HEK293T cells at ∼70–80% confluence were plated in 100-mm dishes and co-transfected with the transfer plasmid (2.6 µg) and Lentiviral Packaging Mix (26 µL; Sigma #SHP001) using X-tremeGENE™ HP DNA Transfection Reagent (Sigma #6366236001) according to the manufacturer’s instructions. Viral supernatants were collected 48 h post-transfection and concentrated using PEG-it® Virus Precipitation Solution (System Biosciences #LV810A-1), then briefly resuspended in cold complete medium. For transduction, target cells were incubated with viral supernatant in the presence of polybrene (7 µg mL⁻¹; Sigma #TR-1003) in complete medium. After 24 h, cells were replated in fresh medium to remove residual virus. Puromycin selection (0.5– 2 µg mL⁻¹; Thermo Fisher Scientific #A1113803) was initiated 48 h post-transduction and maintained for 2–3 days.

### Immunoblotting

Whole-cell lysates were prepared in 1× RIPA buffer (Sigma #20-188) supplemented with 1× Halt™ Protease and Phosphatase Inhibitor Cocktail (Thermo Fisher Scientific #78440). Histones were extracted using the Histone Extraction Kit (Abcam, ab221031) according to the manufacturer’s instructions. Protein concentrations in whole-cell lysates were determined using the Pierce™ BCA Protein Assay Kit (Thermo Fisher Scientific #23228), whereas histone concentrations were measured using the Pierce™ 660 nm Protein Assay (Thermo Fisher Scientific #22662).

Equal amounts of protein from whole-cell lysates or histone extracts were resolved by SDS–PAGE on gels of appropriate acrylamide concentration and transferred onto methanol-activated polyvinylidene difluoride (PVDF) membranes (Thermo Fisher Scientific #88518) by wet transfer in 1× transfer buffer containing 20% methanol at 100 V for 1 h at 4 °C. Membranes were blocked in SuperBlock™ Blocking Buffer (Thermo Fisher Scientific #37535) for 1 h at room temperature and incubated overnight at 4 °C with primary antibodies diluted in blocking buffer. Primary antibodies used were as follows: SLC7A5 (Invitrogen #PA5-115916, 1:1,500); SLC43A2 (Proteintech #30031-1-AP, 1:1,500); Vinculin (Sigma #V9264, 1:2,500); AHCY (Proteintech #10757-2-AP, 1:1,500); ASNS (Cell Signaling Technology #92479, 1:1,000); CAD (Cell Signaling Technology #93925, 1:1,000); GOT2 (Proteintech #14800-1-AP, 1:1,500); H3K4me3 (Cell Signaling Technology #9751, 1:2,000); H3K27me3 (Invitrogen #MA5-11198, 1:1,500); Histone H3 (Cell Signaling Technology #4499, 1:2,000); EZHIP/CXorf67 (Atlas Antibodies #HPA004003, 1:1,000); SLC1A5 (Proteintech #20350-1-AP, 1:1,000); NF-κB p65/RELA (Cell Signaling Technology #8242, 1:1,000); HA tag (Cell Signaling Technology #3724, 1:2,000); ATF4 (Cell Signaling Technology #11815, 1:1,000); phospho-eIF2α Ser51 (Cell Signaling Technology #3398, 1:1,000); and phospho-GCN2 Thr899 (Cell Signaling Technology #94668, 1:1,000). Following primary antibody incubation, membranes were washed three times with 1X Tris-buffered saline (Thermo Fisher Scientific #28358) containing 0.1% Tween 20 (Bio-Rad #1610781) (TBST; 10 min each) and incubated with species-matched infrared secondary antibodies for 1 h at room temperature: IRDye® 800CW donkey anti-rabbit IgG (LI-COR, 926-32213, 1:5,000) or IRDye® 680LT donkey anti-mouse IgG (LI-COR, 926-68022, 1:5,000). Membranes were washed three additional times with TBST, and signals were detected using the LI-COR Odyssey DLx Imaging System.

### Cell proliferation, direct cell counting, and drug-response assays

For longitudinal cell-proliferation assays, cells were seeded in 96-well plates, and cell viability was measured at the indicated time points using PrestoBlue™ Cell Viability Reagent (Thermo Fisher Scientific #P50200) according to the manufacturer’s instructions. Fluorescence was measured at an excitation wavelength of 560 nm and an emission wavelength of 590 nm using a microplate reader. For direct cell counting, cells were harvested, stained with trypan blue, and viable cell numbers were determined using a Countess™ II FL Automated Cell Counter. For drug-response assays, cells were seeded in 96-well plates and treated with serial concentrations of the indicated inhibitors. Cell viability was measured using PrestoBlue™ reagent according to the manufacturer’s instructions. Raw fluorescence values were normalized to the corresponding vehicle-treated controls, and dose–response curves were generated by nonlinear regression in GraphPad Prism 11.1.0 to determine half-maximal effective concentrations (EC₅₀).

### RNA-seq and data analysis

Total RNA was extracted using the Direct-zol™ RNA Miniprep Kit (Zymo Research #R2051) according to the manufacturer’s instructions. RNA concentration, purity, and integrity were assessed using an Agilent 5400 Fragment Analyzer (Agilent Technologies). Poly(A)-enriched, non-directional mRNA libraries were prepared at Novogene (Sacramento, CA). Library quality was evaluated using the Agilent 5400 system, and libraries were sequenced on an Illumina NovaSeq X Plus platform to generate 150-bp paired-end reads, yielding approximately 6 Gb of raw data per sample. Raw sequencing data were processed using Partek® Flow (v10.0; Partek Inc.). Reads underwent pre-alignment quality control, and low-quality bases were trimmed from the 3′ ends using the Trim Bases tool. High-quality reads were aligned to the human reference genome (hg38) using STAR (v2.7.8a) with default parameters and a Partek-built whole-genome index. Gene-level quantification was performed using the Quantify to Annotation Model (E/M) algorithm with Ensembl v109 gene annotation. Low-abundance features were filtered using a noise-reduction threshold ≤ 1. Raw read counts were normalized using the median-ratio method implemented in DESeq2. Differential expression analysis was performed using DESeq2 with default shrinkage estimation. Unless otherwise specified, genes with fold change >1.5, nominal *P* < 0.05, and false discovery rate (FDR) <0.05 were considered significantly differentially expressed. Functional enrichment analyses were performed using the Gene Set Enrichment Analysis (GSEA) desktop application.

### In vitro Limiting Dilution Analysis (LDAs)

LDAs were performed in 96-well plates under 5 or 30 μM methionine conditions. Cells were serially diluted and seeded at 500, 250, 125, or 62.5 cells per well, with eight replicate wells per cell dose. Fresh medium containing the indicated methionine concentration was added every 3 days. Wells were scored for sphere formation, and sphere-initiating cell frequencies were estimated using Extreme Limiting Dilution Analysis (ELDA) software^73^.

### Immunohistochemistry and analysis

Human pediatric ependymoma specimens provided by S.V., A.D., N.K.F., C.-H.G.L., P.S., F.S., E.H., and T.P. were analyzed by immunohistochemistry. RCAS-TVA-derived ependymoma tissue sections were provided by E.H., whereas FFPE blocks and tissue sections from murine IUE-derived ZFTA–RELA fusion-positive ependymomas were provided by T.P. DIPG patient-derived xenograft (PDX) tissues were generated following intracranial implantation of DIPG-13 cells, and medulloblastoma PDX tissues were generated following intracranial implantation of D425 cells. For experimental mouse tissues generated in this study, brains were harvested at experimental endpoints, fixed in 4% formaldehyde for 48 h, transferred to 70% ethanol overnight, and submitted to the University of Pittsburgh Biospecimen Core (PITT) for paraffin embedding, sectioning, and hematoxylin and eosin (H&E) staining. Additional serial unstained sections were generated for immunohistochemical analyses. Formalin-fixed paraffin-embedded (FFPE) tissue sections (5 μm) were deparaffinized in xylene and rehydrated through a graded ethanol series. Antigen retrieval was performed by pressure cooking in citrate buffer (10 mM sodium citrate, 0.05% Tween-20, pH 6.0) for 20 min, followed by quenching of endogenous peroxidase activity using 3% hydrogen peroxide. Sections were blocked in blocking buffer for 30 min at room temperature and incubated overnight at 4 °C with primary antibodies diluted in PBS. Primary antibodies used were as follows: SLC7A5 (Invitrogen #PA5-115916, 1:200); SLC43A2 (Proteintech #30031-1-AP, 1:200); AHCY (Proteintech, 10757-2-AP, 1:200); ASNS (Cell Signaling Technology #92479, 1:200); and SLC1A5 (Proteintech #20350-1-AP, 1:200 and Sigma #HPA035240, 1:200).

Immunoreactivity was detected using the ImmPRESS™ HRP Goat Anti-Rabbit IgG Polymer Detection Kit (Vector Laboratories #MP-7451), according to the manufacturer’s instructions. Signal was visualized using 3,3′-diaminobenzidine (DAB) chromogen (Vector Laboratories #SK-4100). Sections were counterstained with hematoxylin (Fisher Scientific), dehydrated through graded ethanol (70%, 80%, and 100%) and xylene, and mounted using Permount. Representative images were acquired using a bright-field microscope, and staining was quantified from multiple non-overlapping tumor regions per section using ImageJ.

### Chemical compounds and metabolite supplementation and rescue

Small-molecule inhibitors used in this study included the AHCY inhibitor aristeromycin (TargetMol #T10369), the MAT2A inhibitor AGI-24512 (MedChemExpress #HY-112130), the EZH1/2 inhibitor UNC-1999 (MedChemExpress #HY-15646), the DHODH inhibitor BAY-2402234 (MedChemExpress #HY-112645), the SLC1A5 inhibitor V-9302 (MedChemExpress #HY-112683), the G9a/GLP inhibitor UNC0638 (MedChemExpress #HY-15273), the ASNS inhibitor ASX-173 (MedChemExpress #HY-175282), and L-asparaginase (ASNase; MedChemExpress #HY-P1923).

L-Methionine (Thermo Scientific #J61904.18), L-asparagine (Thermo Scientific #371601000), L-aspartic acid (Sigma-Aldrich #A7219), Uridine (Sigma-Aldrich #U3750), and L-homocysteine (Sigma-Aldrich #69453) were used for metabolite supplementation and rescue experiments where indicated.

Stock solutions were prepared in dimethyl sulfoxide (DMSO), sterile phosphate-buffered saline (PBS), or sterile water according to compound solubility and the manufacturer’s recommendations, aliquoted, and stored at −20 °C or −80 °C as appropriate. For in vitro experiments, compounds and metabolites were diluted in culture medium. Vehicle-treated cells received the corresponding concentration of DMSO, PBS, or sterile water.

### Chromatin immunoprecipitation followed by quantitative PCR

Cells were lysed in nucleus preparation buffer [NPB; 10 mM HEPES (pH 7.5), 1.5 mM MgCl_2_, 10 mM KCl, 250 mM sucrose, 0.1% NP-40, and 0.2 mM DTT]^74^. Cell lysates were passed through 100-μm cell strainers and cross-linked with 1% paraformaldehyde (PFA) for 4 min at room temperature with rotation. Cross-linking was quenched by the addition of 125 mM glycine, followed by incubation for 10 min. samples were centrifuged at 1,000*g* for 10 min and the nuclei-containing pellets were washed once in NPB and centrifuged again at 1,000*g* for 10 min. Nuclear pellets were resuspended in nuclei lysis buffer [NLB, 10 mM Tris (pH 8), 1 mM EDTA, 0.1% SDS] and chromatin was sheared using a Qsonica E220 sonicator (Qsonica, Newtown, CT) at 70% amplitude 15 sec sonication pulses followed by 45 sec rest intervals for a total processing time 45 min. Sheared chromatin was centrifuged at 13,000g for 10 min at 4°C to remove insoluble debris and chromatin concentration was determined using a Nanodrop spectrometer.

For chromatin immunoprecipitation (ChiP), 2 μg of sheared chromatin was diluted in ChIP dilution buffer [CDB, 16.7 mM Tris (pH 8), 1.2 mM EDTA, 167 mM NaCl, 1.1% Triton X-100, and 0.01% SDS] and incubated overnight at 4°C with rotation with an antibody against H3K4me3 (1 μg/mL; Cell signaling,) or normal rabbit IgG as a negative control. Protein A/G Dynabeads were washed in PBS containing 1% BSA and subsequently added to the ChIP reactions. Samples were incubated for an additional 2 h at 4°C with rotation. Immunoprecipitated samples were subsequently processed according to the ChIP-IT Express kit protocol (Active Motif, Carlsbad, CA). Beads were collected and washed once with ChIP buffer 1 and three times with ChIP buffer 2 (Active Motif). ChIP DNA was eluted in 100 μl of Elution Buffer AM2 (Active motif) and reverse cross-linked at 65°C for 6 h. The DNA was purified using a MiniElute kit (Qiagen) and eluted in 30 μl of elution buffer. Purified ChIP DNA was analyzed by quantitative PCR (qPCR). ChIP-qPCR was performed using forward primer 5′-GTTTTGCAGTGCGGAGACCA-3′ and reverse primer 5′-TGCAAACTCCACTGGAACCAC-3′, using cycling parameters as previously described^74^.

### ATAC-Seq

ATAC-seq was performed as previously described^18^. Briefly, nuclei were isolated from 50,000 mNSCs transduced with EV or expressing ZFTA–RELA (ZR), with ZR-expressing cells treated with either DMSO or the EZH2 inhibitor GSK343. Nuclei were subjected to Tn5 transposition using the Nextera DNA Library Preparation kit, and amplified libraries were purified and sequenced on an Illumina HiSeq 2500 platform. Reads were trimmed, aligned to the appropriate reference genome, and filtered to remove mitochondrial and duplicate reads. Peaks were identified using MACS2, and BigWig files and enrichment heatmaps were generated using UCSC tools and deepTools, respectively.

### Intracranial tumor implantation

Intracranial tumors were established in C57BL/6J mice by stereotactic implantation of 2 × 10^5^ ZR^Fus^Line_IUE-01 cells into the left cerebral cortex. Equal numbers of male and female mice were included in each experimental group. Before surgery, mice were anesthetized with isoflurane, and meloxicam (5 mg kg⁻¹, intraperitoneally) was administered for perioperative analgesia. The surgical site was disinfected with povidone–iodine and 70% ethanol.

Control and gene-knockdown cells were prepared in sterile phosphate-buffered saline (PBS), and 6 µL of the cell suspension was injected at a depth of 2.5 mm using coordinates of 1 mm left lateral and 1.5 mm anterior to lambda. Cell suspensions were delivered through a burr hole using a Hamilton syringe mounted on a stereotactic apparatus at a rate of 0.6 µL min⁻¹. Following injection, the needle was retained in place for 1 min and then slowly withdrawn to minimize reflux. The incision was closed using wound clips, and mice were monitored until recovery of normal motor activity before being returned to their home cages. Animals were subsequently monitored for postoperative recovery, wound healing, and neurological symptoms. Cohorts implanted with control or gene-knockdown ZR^Fus^Line_IUE-01 cells were used for tumor-burden and survival analyses.

Orthotopic tumors derived from bioluminescent ZFTA–RELA-expressing murine neural stem cells (mNSC_ZR^Fus^) were established in NSG mice by stereotactic implantation of 4 × 10^5^ cells using the same injection volume, coordinates, depth, injection rate, and surgical procedure described above.

### Subcutaneous flank tumor implantation

Subcutaneous tumors were established in female J:NU nude mice by implantation of 5 × 10^5^ bioluminescent ZFTA–RELA-expressing mNSCs into the flank. Tumor establishment was confirmed by bioluminescence imaging (BLI). Once tumors reached a radiance of at least 1 × 10^4^ photons s⁻¹ cm⁻² sr⁻¹, mice were randomly assigned to treatment groups.

### In vivo pharmacological treatments

For the subcutaneous flank tumor study, asparaginase (ASNase; MedChemExpress) was reconstituted in sterile water, aliquoted, and administered at 3 U g⁻¹ body weight by intraperitoneal injection three times per week on alternating days. Vehicle-treated mice received an equivalent volume of sterile water.

For intracranial pharmacological studies, mice bearing mNSC_ZR^Fus^ or ZR^Fus^Line_IUE-01 tumors were randomly assigned to receive vehicle or the dihydroorotate dehydrogenase inhibitor BAY-2402234 (MedChemExpress). BAY-2402234 was formulated in 90% polyethylene glycol 400 (PEG400; MedChemExpress) and 10% ethanol (v/v) and administered at 4 mg kg⁻¹ by oral gavage three times per week on Monday, Wednesday, and Friday for 3 weeks. Vehicle-treated mice received an equivalent volume of 90% PEG400 and 10% ethanol.

### Bioluminescence imaging and analysis

Tumor establishment and longitudinal tumor growth were assessed by BLI using an IVIS Spectrum imaging system (PerkinElmer). Mice were administered D-luciferin (150 mg kg⁻¹, intraperitoneally) and allowed 10 min for substrate distribution before being anesthetized with isoflurane. Animals were maintained under isoflurane anaesthesia throughout image acquisition. Sequential bioluminescence images were acquired to capture the peak signal for each animal. A region of interest was defined over the tumor, and bioluminescence was quantified as radiance, expressed as photons s⁻¹ cm⁻² sr⁻¹. For longitudinal analyses, radiance values were normalized to the corresponding pretreatment baseline value for each animal. In the ASNase flank-tumor study, the day-23 measurement was used as the baseline.

### Survival monitoring and tissue collection

Animals were routinely monitored for general health and neurological symptoms. Mice enrolled in survival studies were euthanized upon reaching predefined humane endpoint criteria, including hunched posture, gait abnormalities, lethargy, or other signs of neurological decline. For the ASNase study, mice were euthanized on day 60, and flank tumors were excised for downstream analyses. For gene-knockdown and BAY-2402234 studies, tissues were collected at matched endpoints when mice in the corresponding control group reached humane endpoint criteria. At these time points, mice from the respective experimental groups were euthanized, and tumors and other tissues of interest were collected for downstream analyses.

### Quantification and statistical analysis

Unless otherwise stated, in vitro experiments were performed with at least three independent replicates. Data are presented as mean ± SD. Statistical analyses were performed using GraphPad Prism 11.1.0. Comparisons between two groups were performed using unpaired two-tailed *t*-tests with Welch’s correction. Comparisons among multiple groups were performed using one-way or two-way ANOVA followed by the indicated multiple-comparisons test (Dunnett’s or Šidák’s). Extreme limiting dilution assays were analyzed using the ELDA likelihood-ratio test. Survival curves were estimated using the Kaplan–Meier method and compared using the log-rank (Mantel– Cox) test. Statistical methods and exact sample sizes (*n*) are indicated in the corresponding figure legends. For transcriptomic and metabolomic analyses, significance thresholds and multiple-testing criteria are specified in the corresponding figure legends. Statistical significance was defined as p < 0.05 (*), p < 0.01 (**), p < 0.001 (***), and p < 0.0001 (****); ns, not significant.

Schematics and graphical illustrations were created using BioRender and GraphPad Prism.

## Notes

### Competing Interest Statement

The authors have declared no competing interest.

