## Extended Data Figures for "Adenosylhomocysteinase drives epigenetic and metabolic reprogramming in ZFTA-RELA ependymoma"

**Extended Data Fig. 1: ZFTA-RELA fusion regulates metabolic programs associated with methionine metabolism.**

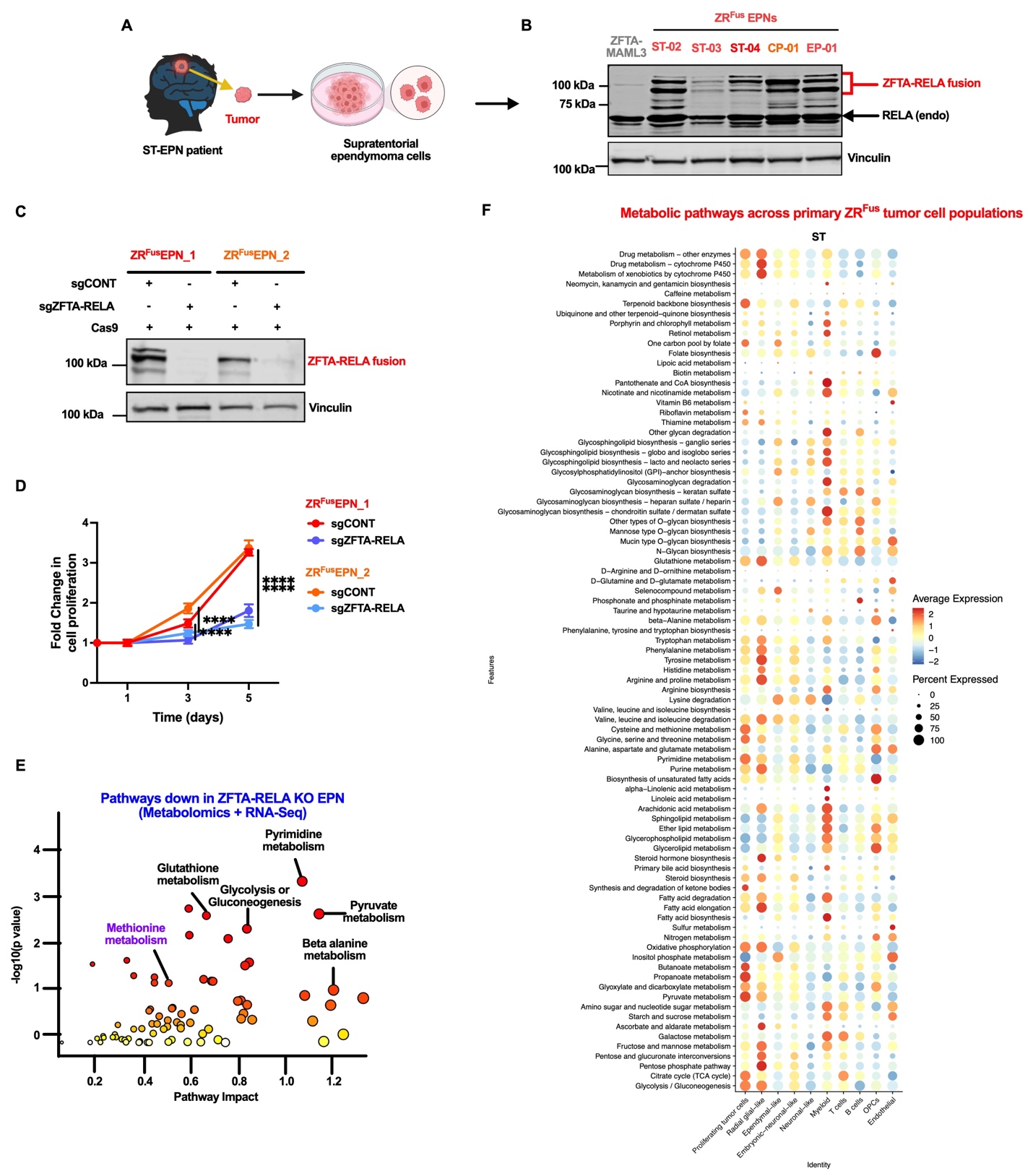

**Extended Data Fig. 2: ZFTA-RELA fusion regulates transcriptional programs associated with methionine metabolism.**

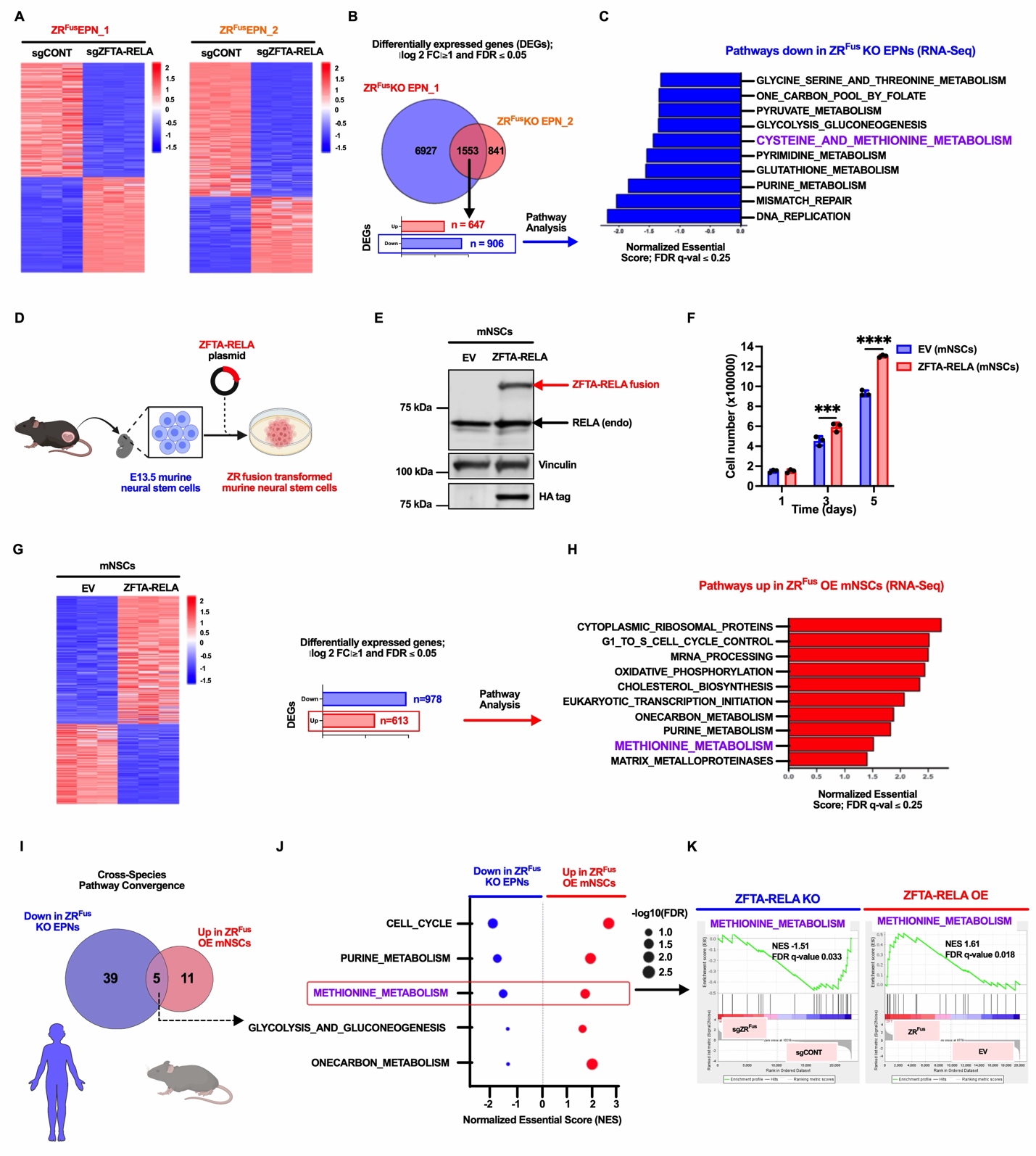

**Extended Data Fig. 3: AHCY regulates tumor growth, methionine-cycle activity, and pyrimidine metabolism in ZFTA-RELA fusion ependymoma.**

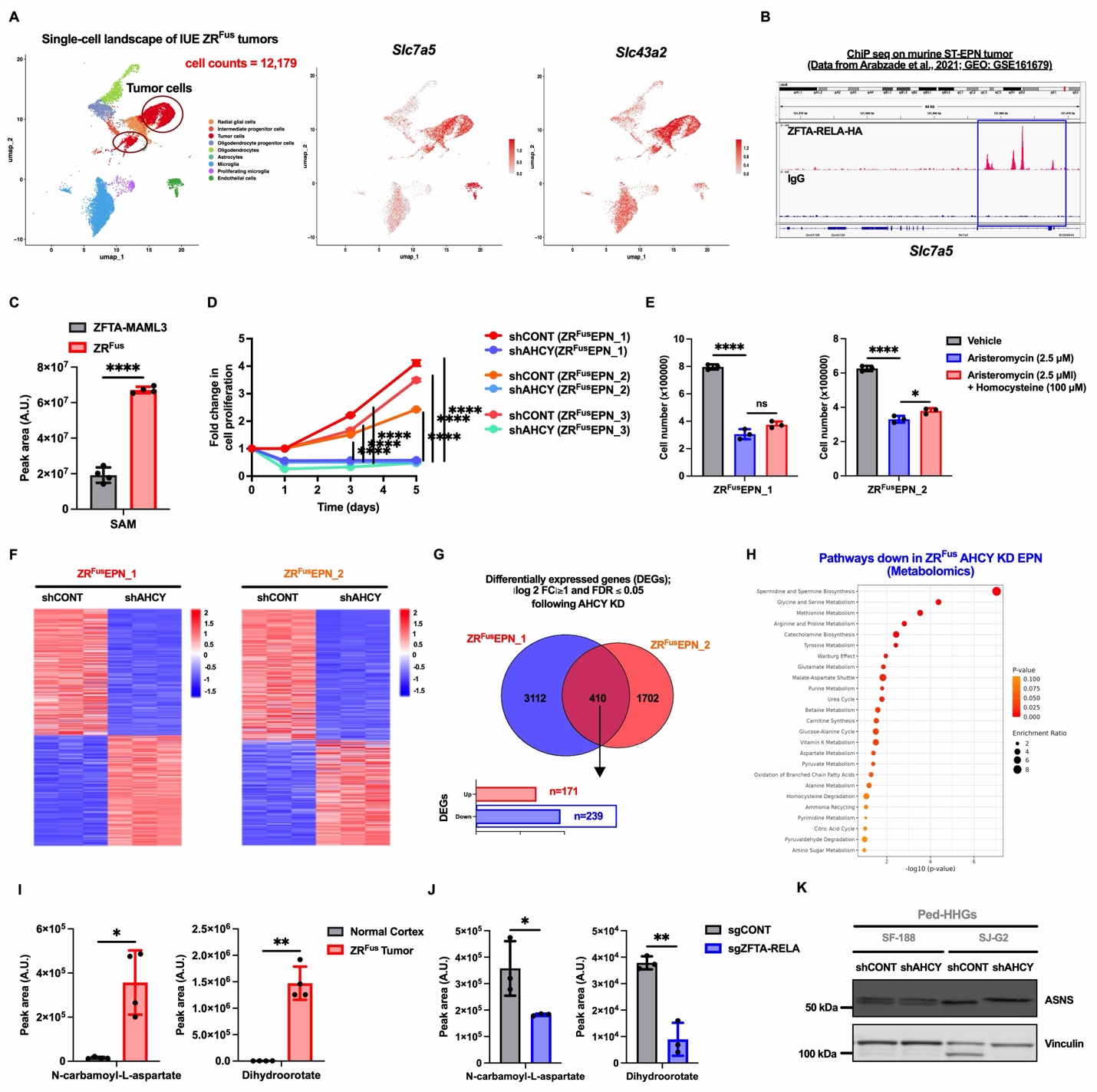

**Extended Data Fig. 4: AHCY regulates ASNS expression independently of the integrated stress response in ZFTA-RELA fusion ependymoma.**

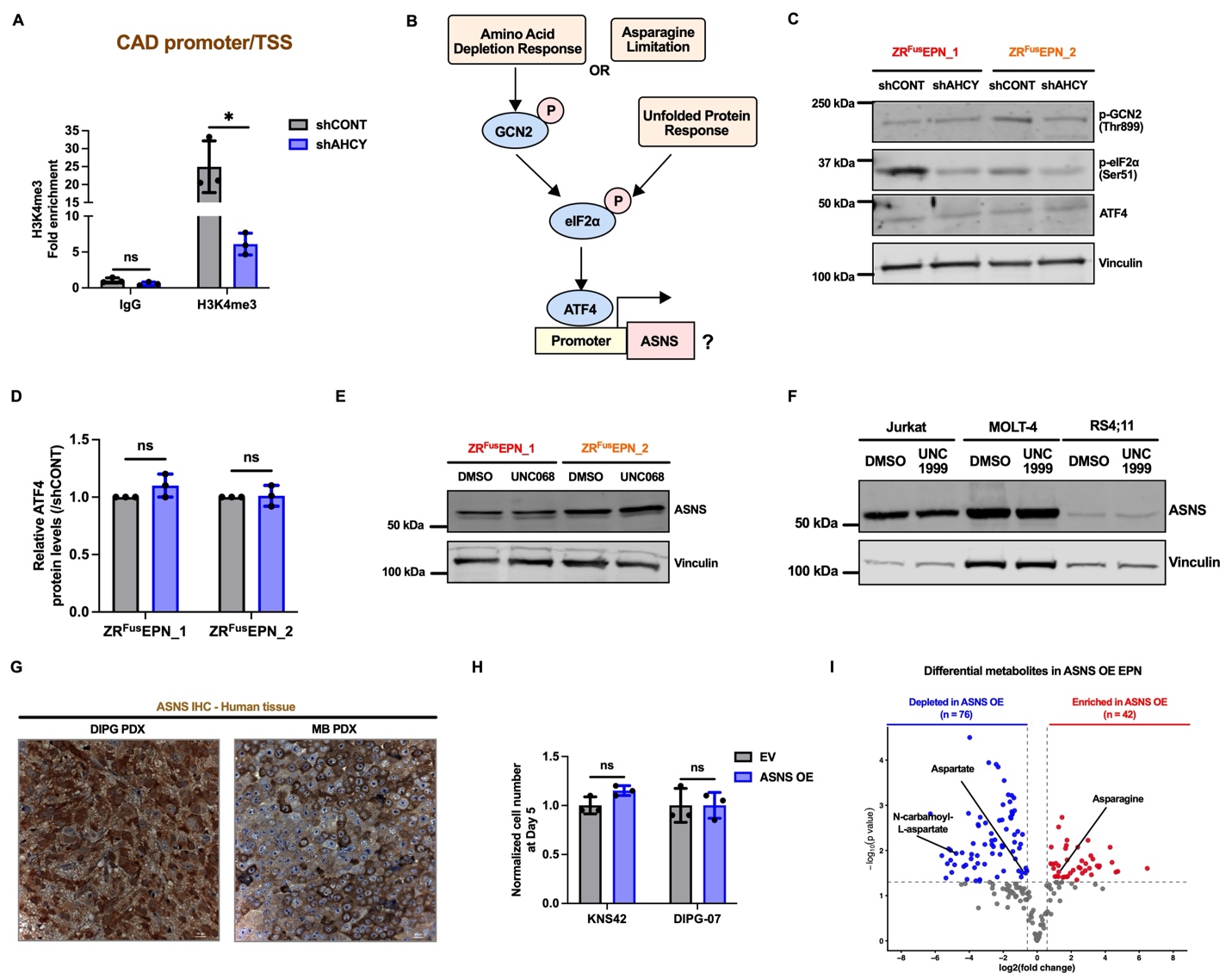

**Extended Data Fig. 5: ZFTA-RELA fusion ependymoma upregulates SLC1A5 expression and is sensitive to asparagine depletion.**

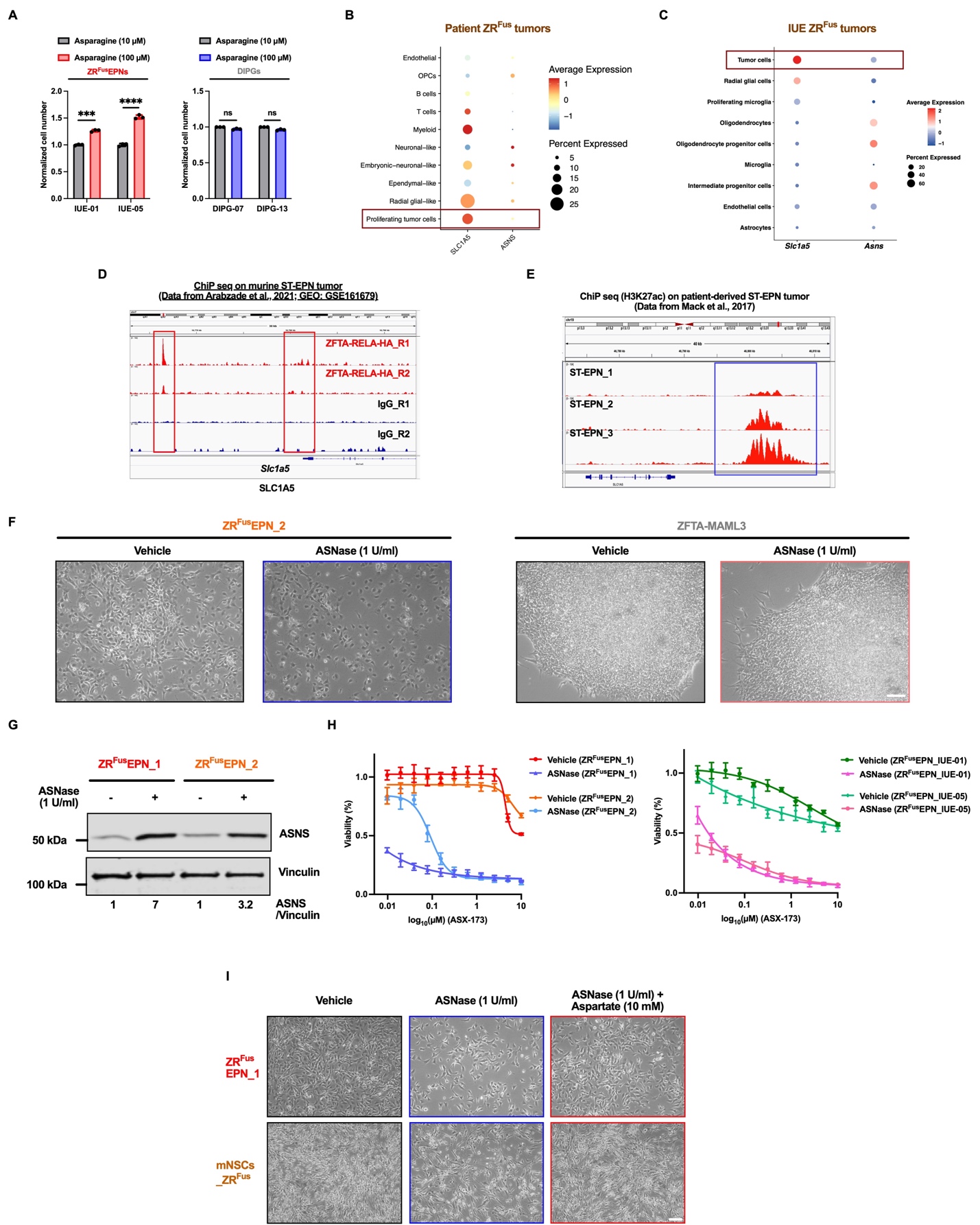

**Extended Data Fig. 6: ZFTA-RELA fusion ependymoma is sensitive to DHODH inhibition.**

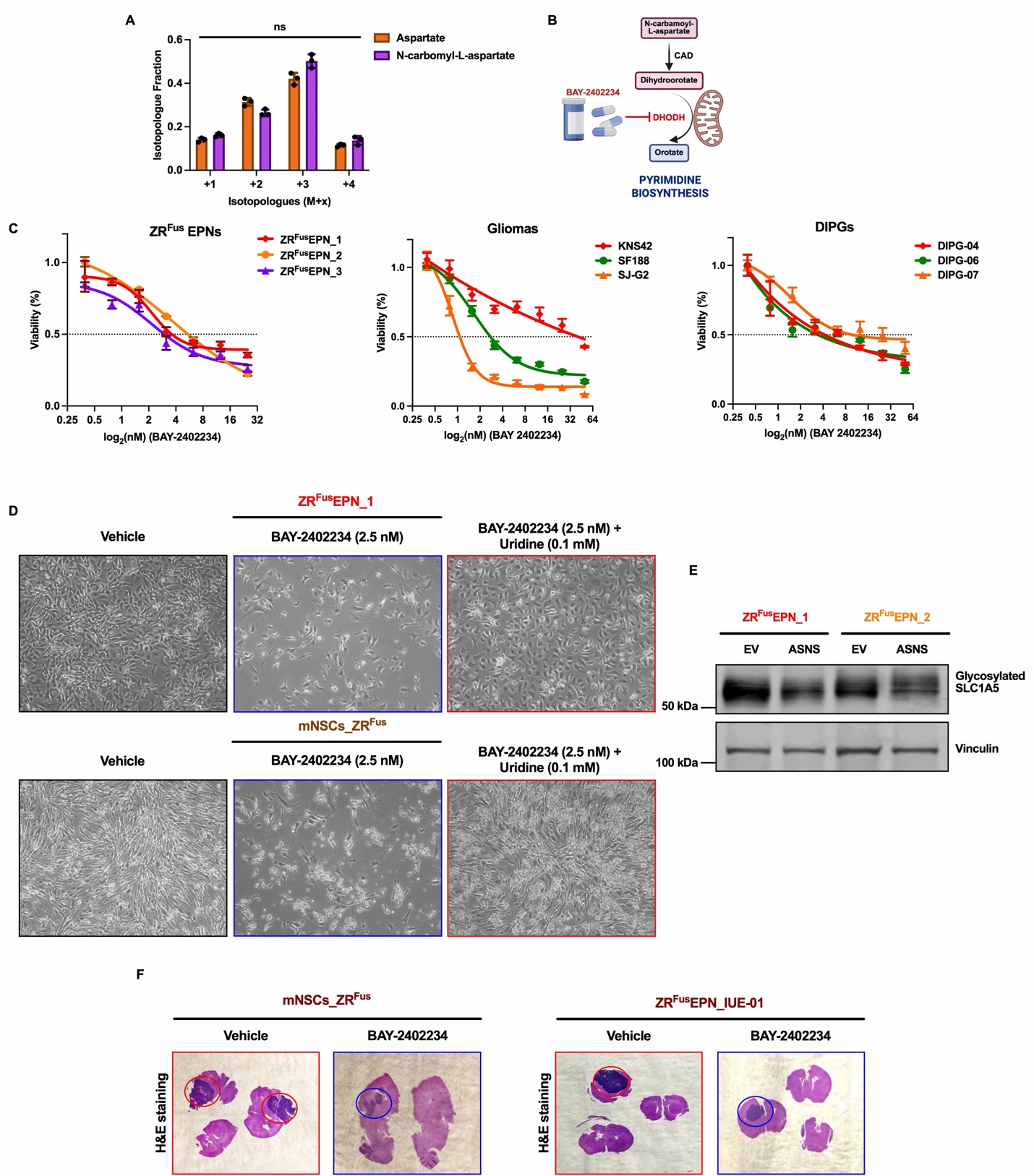
